# Early Partitioning of Structural Paralog Diversity Shapes Immune Evolution Across Habitat Transitions in Gobiiform Fishes

**DOI:** 10.64898/2026.07.27.740993

**Authors:** Katerina L. Zapfe, Ian Birchler De Allende, Aditt Mahadik, Gerard R. Nasser, Bruno Frédérich, Jeffrey A. Yoder, Alex Dornburg

**Affiliations:** Department of Bioinformatics and Genomics, University of North Carolina at Charlotte, Charlotte, North Carolina, USA; Department of Biological Sciences, North Carolina State University, Raleigh, North Carolina, USA; Genetics and Genomics Academy, North Carolina State University, Raleigh, North Carolina, USA; Laboratory of Evolutionary Ecology, FOCUS, University of Liège, Liège, Belgium; Department of Molecular Biomedical Sciences, College of Veterinary Medicine, North Carolina State University, Raleigh, North Carolina, USA; Comparative Medicine Institute, North Carolina State University, Raleigh, North Carolina, USA

**Keywords:** Innate immunity, gene family evolution, paralog diversification, ecological transitions, protein structural evolution, comparative genomics, immunogenetics

## Abstract

Habitat transitions expose species lineages to novel pathogen regimes and are often hypothesized to drive adaptive diversification of immune gene families. However, the temporal association between gene family diversification and ecological change remains unresolved. To gain deeper insight into the relationship between the diversification of species and the evolution of their immune system, we investigated the evolutionary history of Toll-like receptors (TLRs) across Gobiiformes, a clade characterized by repeated transitions across aquatic and amphibious environments. TLRs are well-studied membrane-bound pattern recognition receptors that play crucial roles in detecting pathogens and immune activation. Phylogenomic, structural, and sequence analyses reveal that a major expansion of TLR22 predates many ecological transitions, with early paralog diversification partitioning receptor architectures into distinct structural regimes that persist across lineages. Subsequent evolution is concentrated in the extracellular ligand-binding, leucine-rich repeat (LRR) domains, where localized sequence and structural variation enables likely functional tuning without major architectural innovation. These results indicate that ecological transitions do not require repeated evolution of new immune receptor forms, but can instead be facilitated by reconfiguration of pre-existing immunogenetic diversity. These findings raise the possibility that expansions of immune gene families occurring early in a clade’s evolutionary history may commonly persist and subsequently act as substrates for evolutionary responses to environmental change.

## Introduction

Evolutionary habitat transitions are fundamental aspects of the Tree of Life, spanning transitions from terrestrial to aquatic environments (Dunlop et al. 2013; Vermeij and Motani 2018), fresh to marine waters (Carrete Vega and Wiens 2012; Betancur-R et al. 2015), shallow to deep waters (Rincon-Sandoval et al. 2020; Anger and Urzúa 2024), or surface to subterranean environments (Culver and Pipan 2019; Recknagel and Trontelj 2022) to name but a few. Such transitions often coincide with dramatic changes in physiology (Lee et al. 2003), morphology (Farisenkov et al. 2022; Zhao et al. 2024), or behavior (Chapman et al. 2012) that in turn can enable organisms to radiate in response to new ecological opportunities (Stroud and Losos 2016). Recent comparative genomic investigations have begun to reveal the molecular mechanisms underlying these habitat transitions (Aristide and Fernández 2023), frequently implicating multigene families (groups of related genes arising from duplication events that diversify to perform specialized functions) as critical contributors to successful ecological shifts (Wang et al. 2016; O’Connor et al. 2018; Waters and Vierling 2020; Dornburg et al. 2022; Akiyama et al. 2022). However, the temporal relationship between diversification in these gene families and the ecological transitions with which they are associated is often unresolved. As a result, it is unclear whether such genomic changes commonly represent standing diversity that facilitated the initial transition, emerged concurrently with it, or represent subsequent diversification.

The gene families that form the molecular basis of the innate immune response represent a primary interface between organisms and their environments. Habitat shifts expose lineages to novel microbial communities and altered pathogen regimes, and the ability of an individual to mount a response through the recognition of conserved pathogen-associated molecular patterns (PAMPs) forms a front line of defense that is especially critical when confronted with unfamiliar pathogens (McDade et al. 2016). Accordingly, variation in innate immune receptors has been linked to ecological performance in novel environments (Becker et al. 2020). For example, elevated expression of specific innate immune receptors has been observed in range-edge populations of house sparrows, suggesting a potential role for immune responsiveness in mediating invasion dynamics (Martin et al. 2014, 2017). However, at deeper evolutionary timescales, the diversity of multigene families is often shaped by the birth and death of paralogs, resulting in highly uneven distributions of paralog richness across species for the same gene family (Niimura 2009; Liu et al. 2020; Wcisel et al. 2023). This asymmetry raises a question: when does the functional diversity associated with these paralogs arise relative to ecological transitions? One possibility is that functional diversification between paralogs occurs prior to ecological change, with ancient gene duplications retained over long timescales preserving structurally differentiated recognition capacity that can later be deployed in new environments. Alternatively, functional diversification may occur during ecological transitions, with episodic bursts of duplication and divergence generating novel receptor structure and ligand-binding properties in response to new pathogen regimes. Finally, functional diversification may largely occur after ecological transitions, reflecting ongoing host–pathogen coevolution that reshapes immune repertoires following establishment in new environments. Disentangling these temporal dynamics is essential for determining whether immune repertoires associated with major habitat shifts reflect pre-existing genomic substrates, contemporaneous innovation, or subsequent coevolutionary remodeling.

Toll-like receptors (TLRs) are among the most well-studied multigene families of innate immune receptors, and serve as a model for understanding the molecular evolution of the immune response (Velová et al. 2018; Liu et al. 2020; Carlson et al. 2022; Boraschi et al. 2023). As pattern recognition receptors (PRRs), TLRs initiate innate immune responses following the detection of PAMPs or endogenous damage-associated molecular patterns (DAMPs) (Tang et al. 2012). Functional studies have demonstrated that different TLRs mediate defense against diverse classes of pathogens, including fungal, bacterial, viral, and parasitic infections (Netea et al. 2007; Gerold et al. 2007; Zhang et al. 2014; Fouzder et al. 2020; Kayesh et al. 2021), establishing a direct link between receptor diversity and immune function. Comparative genomic analyses have revealed that the diversity of TLRs that underlies this defense is markedly uneven across vertebrates (Velová et al. 2018; Khan et al. 2019; Liu et al. 2020; Stejskalova et al. 2024). Extensive lineage specific paralog expansions have been identified and proposed to provide an evolutionary advantage to pathogen defense by expanding the repertoire of detectable ligands through the formation of heterodimers (Solbakken et al. 2016; Khan et al. 2019). However, TLR diversity is expressed not only through variation in paralog number, but also through diversification of receptor architecture. The extracellular leucine-rich repeat (LRR) domain forms a conserved structural scaffold whose repeat number, three-dimensional organization, and surface topology collectively determine ligand recognition and receptor dimerization, allowing relatively modest structural modifications to alter ligand specificity while preserving downstream signaling (Botos et al. 2011; Bzówka et al. 2023; Chen et al. 2024). Indeed, signatures of selection in ligand-binding LRR domains within these expansions have been used to suggest increased recognition breadth may be advantageous for lineages encountering diverse microbial communities during ecological transitions (Solbakken et al. 2017; Liu et al. 2020; Włodarczyk et al. 2023). Therefore, understanding how ecological transitions shape immune evolution requires determining not only when new paralogs originate, but also resolution of the evolutionary dynamics of TLR structural changes within a comparative system that integrates phylogenetic context with well-characterized habitat shifts.

Gobiiform fishes (order Gobiiformes containing Apogonoidei, Gobioidei and *Trichonotus sensu stricto*, Near and Thacker (2024)) provide a powerful comparative system for investigating the temporal dynamics of immune gene family diversification surrounding ecological transitions. Comprising over 2000 species, gobiiforms are characterized by the repeated colonization of diverse habitats that span marine, estuarine, freshwater, subterranean, and amphibious environments (Thacker 2009, 2015; de Brito et al. 2022; McCraney et al. 2025). Among these, the mudskippers are widely regarded as living analogs of early tetrapods due to their capacity for sustained terrestrial activity and the morphological and physiological innovations that enable life at the water-land interface (You et al. 2018). Genome sequencing has revealed striking expansions of TLR paralogs in multiple mudskipper species, which have been hypothesized to reflect expansion in response to novel infection pressures imposed by terrestrial pathogens (Qiu et al. 2019). However, similar TLR expansions have also been identified in the distantly related round goby (*Neogobius melanostomus*), an ecologically versatile species capable of rapidly invading new areas (Adrian-Kalchhauser et al. 2020). Whether these TLR expansions in two distantly related gobiiform species reflect ancient expansions that predate habitat transitions, independent bursts of functional immunogenetic diversification associated with ecological shifts, or subsequent lineage-specific remodeling following establishment in new environments remains unknown. The scale and repetition of habitat transitions across gobiiforms provide an unprecedented opportunity to resolve these alternatives and temporally contextualize immunogenetic implications of crossing ecological boundaries over macroevolutionary timescales.

Here, we integrate analyses of publically available transcriptomes and genomes with a comparative phylogenomic approach to assess the diversification dynamics of TLR paralogs across the Gobiiform radiation. Leveraging this comparative framework, we first quantify patterns of paralog evolution and identify a dramatic, previously unknown paralog expansion in the early evolutionary history of Gobiiformes. We then assess whether variation in paralog richness is associated with habitat transitions by reconstructing ancestral TLR repertoires, revealing that these expansions in mudskippers substantially predate the origin of amphibious life histories. To evaluate whether this standing molecular diversity corresponds to functional divergence, we modeled the protein shape of each paralog using artificial intelligence based models of protein folding. By embedding these models within a phylomorphospace, we quantified structural divergence across paralogs and lineages, revealing that much of the TLR functional landscape was already in place prior to previously suggested ecological drivers of immunogenetic diversification. These findings suggest that ancient expansions in TLR functional repertoires have acted as a scaffold for ecological innovation, offering new insight into how pre-existing immunogenetic diversity shapes evolutionary potential during major habitat transitions.

## Materials and Methods

### Data Acquisition

We selected species that span the backbone of the gobiiform phylogeny from available public genomic and transcriptomic resources on NCBI. As chromosome-level high coverage reference genomes remain limited, the addition of transcriptomic data allowed us to survey TLR diversity across a greatly expanded pool of species (**Supplemental Table S1**). TLRs have a high baseline level of expression in various tissues including spleen, gill, heart, and liver among others in fishes (Thompson et al. 2021; Zhang et al. 2022; Liu et al. 2024), rendering comparative transcriptomic investigations of TLRs a common alternative approach to reliably assessing their diversity in the absence of genomic sequence data (Tong et al. 2015; Eggestøl et al. 2018).

Genome databases were searched using BLAST (Sayers et al. 2021) with TLRs previously identified from *Boleophthalmus pectinirostris*, *Periophthalmus magnuspinnatus*, *Sphaeramia orbicularis*, and *Eucyclogobius newberryi* as search queries (Qiu et al. 2019; Carlson et al. 2023). Candidate sequences were retained using an e-value threshold below e-10. For transcriptomic datasets, only transcripts derived from tissues with known high TLR baseline expression (e.g., gill, liver, and spleen) were used. Transcriptomes derived from a mix of tissues, such as “liver, heart, gut, ovary, brain, limbs, skin…” were included if at least one of the three target organs was present. For experimental studies with varying treatments, such as salinity stress, only control group sequences were included to avoid conditions potentially leading to a down regulation of TLR expression. Collectively, the genome and transcriptome search identified sequence data from a total of 27 Gobiiformes: 16 species of Gobiidae, 4 mudskipper species from the genera *Boleophthalmus, Periophthalmodon,* and *Periophthalmus*), 2 species from Butidae, 1 species from Odontobutidae, and 4 species from Apogonidae (**Supplemental Datasheet 1**).

FASTQ files for each species were retrieved from the NCBI SRA database using SRAtools (v3.1.0) with no limit on the maximum file size. Transcriptomes were assembled using Trinity v2.15.1 (Grabherr et al. 2011). Transcript completeness was assessed using BUSCO v5.4.7 to identify ultraconserved, single copy genes within our transcriptome assemblies (Simão et al. 2015) using the Actinopterygii ortholog database version odb10 (Zdobnov et al. 2017). For species with multiple candidate transcriptome datasets, only the assembly with the highest BUSCO score was retained for primary analyses (**Supplemental Fig. S1**). However, the remaining assemblies were used to assess the sensitivity of sequence searches to available transcriptomic data (see “*Reconstructing the ancestral history of paralog duplication”* below). Open reading frames (ORFs) were identified using the TransDecoder.LongOrfs function in TransDecoder v5.7.1 (https://github.com/TransDecoder) with candidate coding regions predicted using the *Predict* function.

Candidate TLR protein sequences were identified from TransDecoder-generated peptide files (.pep) using the hmmsearch function in HMMER (http://hmmer.org/). This search was based on a curated guide FASTA file that included all acanthomorph TLRs identified by Carlson et al. (2023). Putative TLRs were extracted using the HMMParse function in the TOAST R-package (v4.4.2, (Wcisel et al. 2020), retaining only hits with e-values less than 1e-10. To validate domain architecture, extracted sequences were screened against the PFAM database (Bateman et al. 2000) in conjunction with the HMMScan function in HMMER. Sequences containing domains inconsistent with TLRs such as death or immunoglobulin domains were excluded. For retained sequences, Toll/interleukin-1 receptor (TIR) and leucine-rich repeat (LRR) domains were identified using the HMMScanParse function in TOAST (Wcisel et al. 2020).

Following identification of taxa with immunogenetic data, we additionally collected data on habitat (marine, freshwater, or brackish) as well as life history (amphibious Vs aquatic). For each taxon, all data was collected from Fishbase (Pauly 2010) using the rFishbase package (Boettiger et al. 2012) (**Supplemental Table S1**).

### Phylogenetic Verification and Nomenclature of TLR Sequences

To evaluate the diversity of TLR sequences, we used the TLR classification framework from Carlson et al. (2023), which integrated phylogenetic inference with naming conventions to standardize vertebrate Toll-like receptor (TLR) nomenclature. In this framework, TLR sequences are assigned names relative to well-characterized reference genes from multiple taxa, including humans (Tweedie et al. 2021), zebrafish (Bradford et al. 2022), and bowfin (Thompson et al. 2021), based on their resolution within a robust phylogenetic framework. This approach addresses widespread inconsistencies stemming from automated gene annotations for TLRs in ray-finned fishes which misidentified at least 40% of the species surveyed by Carlson et al. (2023). Accordingly, we reassigned TLR names based on the phylogenetic placement of sequences within these well-established clades rather than rely on existing annotations alone for determining TLR identity. For example, if a sequence originally annotated as TLR4 was to be resolved as belonging with the TLR18 clade, it would be reclassified as TLR18. For closely related paralogs within a clade, genus-level or species-level identifiers were appended to distinguish copies. All TLR sequences identified in this study were named following this framework, thereby aligning our results towards a unified framework for understanding vertebrate TLR diversity.

Phylogenetic analyses were conducted using TIR domain sequence identified from gobiiform transcripts and genomes were aligned together with previously identified ray-finned fish TLR TIR domains (Carlson et al. 2023). This allowed us to place gobiiform TLRs into the broader phylogenetic context of their affinity to previously identified and named TLR lineages. The best-fitting model of amino acid substitution was selected using ModelFinder (Kalyaanamoorthy et al. 2017) as implemented in IQ-TREE v2 (Minh et al. 2020). ModelFinder evaluates a broad candidate pool of amino acid substitution models that include not only commonly used models such as JTT and WAG, but also incorporates the possibility of empirical profile mixture models, and several options for accommodating rate heterogeneity (e.g., discrete gamma and FreeRate models) (Wong et al. 2025). Maximum likelihood based phylogenetic inference was conducted using IQ-TREE v2 (Minh et al. 2020), with node support assessed using 1000 ultrafast bootstrap replicates (Hoang et al. 2018).

Restricting our phylogenetic analyses to TIR domains minimized alignment ambiguity and the effects of substitution saturation [e.g., phylogenetic noise (Townsend et al. 2012; Dornburg et al. 2017, 2019, 2026)]. Toll-like receptors exhibit domain specific evolutionary dynamics associated with their role in recognizing PAMPs and initiating downstream immune signalling. The TIR domain, which mediates signal transduction (Valkov et al. 2011), is typically more conserved and retains phylogenetically informative signal across distantly related taxa (Roach et al. 2005; Carlson et al. 2023). However, The extracellular LRR domain, which mediates ligand recognition and binding (Leonard et al. 2009), evolves rapidly resulting in high levels of alignment uncertainty at deep evolutionary timescales due to high sequence divergence (Roach et al. 2005; Mikami et al. 2012; Carlson et al. 2023).

### Reconstructing the ancestral history of paralog duplication

We reconstructed the evolutionary history of TLR paralog diversification across Gobiiformes, by building upon a previously published phylogenomic dataset comprising 1,314 ultraconserved element (UCE) loci sampled from 121 species (McCraney et al. 2025). This dataset comprises representatives of all major gobiiform lineages (McCraney et al. 2025), as well as several outgroups that span the backbone of early acanthomorph divergences (Near et al. 2012; Hughes et al. 2018; Dornburg and Near 2021). We expanded taxonomic coverage to incorporate additional species represented in our transcriptomic data but absent from the UCE matrix by supplementing this dataset with sequences from four commonly used loci [*12S*, *cytochrome b (CYTB)*, *COI*, and *RAG1*] retrieved from GenBank (**Supplemental Table S2**). UCE and Sanger-derived loci were concatenated using TOAST (Wcisel et al. 2020), with loci treated as candidate partitions for phylogenetic inference. As highly non-random missing data patterns are known to promote spurious topological inference (Simmons 2012), we placed the additional species within their respective genera using functions from the RRPhylo R-package (Castiglione et al. 2018) and constraining the topology to follow the backbone based on the UCE loci. Partition and model selection were performed using ModelFinder (Kalyaanamoorthy et al. 2017) as implemented in IQ-TREE v2 (Minh et al. 2020). Following model selection, branch lengths were optimized on the constraint topology IQ-TREE v2 (Minh et al. 2020). This strategy allowed us to leverage the power of the UCE loci towards branch length estimation while mitigating possible errors due to substitution saturation that is common in mitochondrial loci at these timescales (Dornburg et al. 2014, 2024).

To place the inferred phylogeny into a temporal framework, we estimated divergence times from the maximum likelihood tree using the RelTime method (Tamura et al. 2012). RelTime estimates node ages under a relative rate framework that relaxes the assumption of a strict molecular clock as well as assumptions of rate autocorrelation/independence (Tamura et al. 2018). Instead divergence times are directly computed from branch lengths and scaled using calibration constraints (Tamura et al. 2018), yielded estimates comparable to those from Bayesian methods while maintaining computational efficiency and robustness to between lineage rate heterogeneity (Battistuzzi et al. 2018), a possible major source of error (Dornburg et al. 2012) that may be prevalent at deep time scales (Berv et al. 2024). We implemented RelTime in MEGA v12 (Kumar et al. 2024) using the divergence times estimated by McCraney et al. (2025) as secondary calibrations (**Supplemental Table S3**) that were used to set mean ages, minimum, and maximum bounds at calibrated nodes (Tao et al. 2020), setting the clade containing *Lampris, Polymixia,* and *Percopsiformes* as the outgroup (Dornburg and Near 2021; Ghezelayagh et al. 2022; McCraney et al. 2025). Following time calibration, we used the *Phytools* R-package v2.0, (Revell 2024) to map variation in TLR copy numbers onto the gobiiformes phylogeny. Ancestral state reconstructions of paralog counts over time were assessed using the phenogram function which estimates ancestral states using the *fastAnc* function, allowing us to quantify the degree to which expansions TLR paralog richness align with their evolutionary transition out of a purely aquatic lifestyle. To assess the sensitivity of ancestral reconstructions to incomplete or fragmented transcriptomes, which may limit detection of full-length paralogs or inflate gene counts through splice variants, we repeated all analyses using additional transcriptomic datasets derived from multiple individuals per species obtained from GenBank. This sensitivity analysis allowed us to evaluate the robustness of inferred ancestral states to variation in transcript representation and sequencing completeness.

### TLR structural evolution

To evaluate how structural diversification of TLR proteins unfolds through evolutionary time, we constructed a protein “phylomorphospace” (Sidlauskas 2008) that captures variation in three-dimensional geometry across paralogs and lineages. This framework extends concepts from landmark-based geometric morphometrics (Zelditch et al. 2012) to protein structure, allowing us to assess whether expansions in TLR copy number are associated with expansions in structural space. To this end, we curated a set of full-length amino acid sequences that included the expected complement of LRR domains and a single TIR domain. Domain composition initially predicted from HMMER was verified using SMART (Simple Modular Architecture Research Tool; https://smart.embl.de/) to ensure completeness and correct annotation (Schultz et al. 2000). Sequences lacking required domains, exhibiting truncated architectures, or showing evidence of incomplete annotation were excluded, yielding a dataset with amino acid sequences associated with protein structures for 27 species (**Supplemental Datasheet 2**).

Three-dimensional protein structures of the validated sequences were predicted using ESMFold (Lin et al. 2023). From each predicted structure, α-carbon (Cα) coordinates were extracted to represent the geometric backbone of the protein. To establish residue correspondence across structures, we generated multiple sequence alignments using FAMSA for the full TLR dataset and TLR subclades of interest. Alignment columns were filtered to retain only positions with complete coverage across all sequences, ensuring that each structure contributed an identical set of homologous residues and that we maintained a consistent coordinate framework for geometric comparisons.

Filtered Cα coordinate sets were superimposed using Generalized Procrustes Analysis (GPA) to remove differences due to translation, rotation, and scaling, while preserving the spatial relationships among residues (Zelditch et al. 2012) using custom code in Python. For each aligned structure, we calculated all pairwise Euclidean distances between homologous Cα atoms, generating a distance-based representation of internal fold geometry as the basis for multivariate analyses of structural variation. Using the *phyl.pca* function from phytools v2.0 (Revell 2024) we integrated this representation of the internal fold morphology with the maximum likelihood estimated Phylogeny of paralog divergences based on TIR domains to quantify the major axes of protein shape variation while accounting for shared evolutionary history. Resulting ordinations were visualized as phylomorphospaces (Sidlauskas 2008), using the phylomorphospace function from phytools v2.0 (Revell 2024). To assess whether evolutionary habitat transitions such as those experienced by mudskippers promote dispersion in morphospace occupancy, we conducted a permutational multivariate analysis of variance (PERMANOVA) with 999 permutations (Anderson 2017), using the the adonis2 function in the R-package vegan v.2.7 (Dixon 2003).

To assess whether our results were robust to alternate representations of protein structure, we employed the FAIR-ESM transformer-based neural network (Rives et al. 2021) to extract high-dimensional structural feature embeddings from predicted protein models. These embeddings were again summarized using phylogenetic PCA (Revell 2009) to visualize fine-scale patterns of structural differentiation within and among Gobiiformes. Together, these analyses provide a complementary view of structural diversification, integrating explicit geometric comparisons with learned representations of protein structure.

### Sequence variation in the recognition architecture of TLRs

We analyzed the variability of LRR domains extracted from aligned TLR sequences to characterize patterns of sequence conservation and variability within the domain associated with TLR ligand recognition/binding function. TLRs generally possess multiple LRR domains within the extracellular region. To visually assess whether variation in domain frequencies between sequences reflects LRR expansions within core domain containing segments, we plotted the architecture of the LRR domains across each sequence using ggplot. We then tested the relationship between LRR expansions, contractions, and intervening extracellular sequences in TLRs, using the number and length of LRR repeats for each sequence using domain annotations from HMMER and SMART. We performed a phylogenetic least-squares regression of total sequence length against LRR count, combined LRR length, and the total length of non-LRR extracellular regions, as each is expected to scale with overall protein size. This approach allowed us to isolate variation in domain architecture independent of size and extract residuals from this model that were subsequently used in a series of phylogenetic generalized least squares regressions using a Brownian motion (BM) model (Grafen 1989; Martins and Hansen 1997) to evaluate the relationships among extracellular domain length, LRR number, and total LRR length. These analyses allowed us to test whether variation in extracellular domain size is primarily driven by LRR expansion or whether such expansions require coordinated changes in intervening sequences.

To place patterns of domain expansions or contractions into an evolutionary context, we mapped LRR number and total LRR length onto the paralog phylogeny and quantified the associations between LRR number and total LRR length with phylogenetic relationships using tests of phylogenetic signal. We first assessed Pagel’s λ, which provides a measure from 0 (phylogenetic independence) to 1 (phylogenetic dependence) on the degree to which trait variation is consistent with a BM model of evolution. We then utilized Blomberg’s K as a complementary approach to detect deviations from BM such as overdispersion or convergence. Both statistics were implemented using the phylosig function in the R package phytools (Revell 2024). We additionally estimated the evolution of LRR domain copy number across the paralog phylogeny through ancestral state analyses using the *fastAnc* function in phytools.

Finally, we tested whether lineages belonging to the expanded TLR clades exhibited different average rates of LRR domain expansion and contraction than the remaining sampled TLR paralogs using evolutionary rates estimated with BAMM (Rabosky 2014). Although BAMM is commonly used to identify discrete evolutionary rate shifts across phylogenies (Ghezelayagh et al. 2022; Dimitrov et al. 2023; Dornburg et al. 2024), we did not employ a formal shift-detection framework because our paralog phylogeny was assembled to compare focal TLR clades with a representative background of sampled paralogs rather than to infer evolutionary rate regimes across a comprehensively sampled phylogeny of all gobiiform TLRs. Instead, we compared the posterior distributions of branch-specific evolutionary rates between focal and background TLR lineages, providing a direct assessment of whether the former occupy a distinct region of LRR evolutionary rate space within the sampled phylogeny. LRR domain number was treated as a continuous character and priors were estimated using BAMMtools (Rabosky et al. 2014). Markov chain Monte Carlo analyses were run for 50 million generations, sampling every 1,000 generations. Convergence and mixing were evaluated by inspecting likelihood traces and confirming adequate effective sample sizes (ESS) for all estimated parameters.

## Results and Discussion

### Paralog expansions occurred early in Gobioid evolutionary history

Our analyses recovered 202 unique TLR sequences across gobiiforms, all of which resolve within well established TLR clades (**Fig. 1, Supplemental Datasheet 1** & **Supplemental Figs. S2-S14**) and are consistent with previously described patterns of TLR family presence and absence in acanthomorph fishes (Carlson et al. 2023). For example, TLR6 and TLR10 are likely mammalian-specific and thus absent from all Actinoptergii (Huang et al. 2011; Mikacenic et al. 2013; Carlson et al. 2023) and, with the exception of soldierfish (*Myripristis*), TLR4 has not been reported in other acanthomorphs (Carlson et al. 2023). TLR22 represents a pronounced exception to this pattern, exhibiting substantial expansion in copy number that mirrors paralog expansions reported in genome-centric studies of mudskippers (You et al. 2018; Kim et al. 2021) and round gobies (Adrian-Kalchhauser et al. 2020) (**Figs. 1-2**). However, Ancestral reconstruction places the increase in copy number deeper within Gobioidei, indicating that both mudskippers and round gobies inherit an already expanded receptor repertoire (**Fig. 3A & Supplemental Fig. S15**). Rather than being restricted to amphibious or invasion-prone taxa (**Fig. 3B**), expanded TLR22 repertoires are distributed across multiple gobioid clades (**Fig. 3C**). These results are consistent across replicate transcriptomic datasets, identifying the TLR22 expansion as a clade-wide feature of gobioid evolution. Collectively, these results suggest that many lineages have long maintained elevated numbers of TLR22 sequences as a standing substrate for pathogen defense across diverse aquatic environments (**Fig. 3C**).

**Fig. 1.**
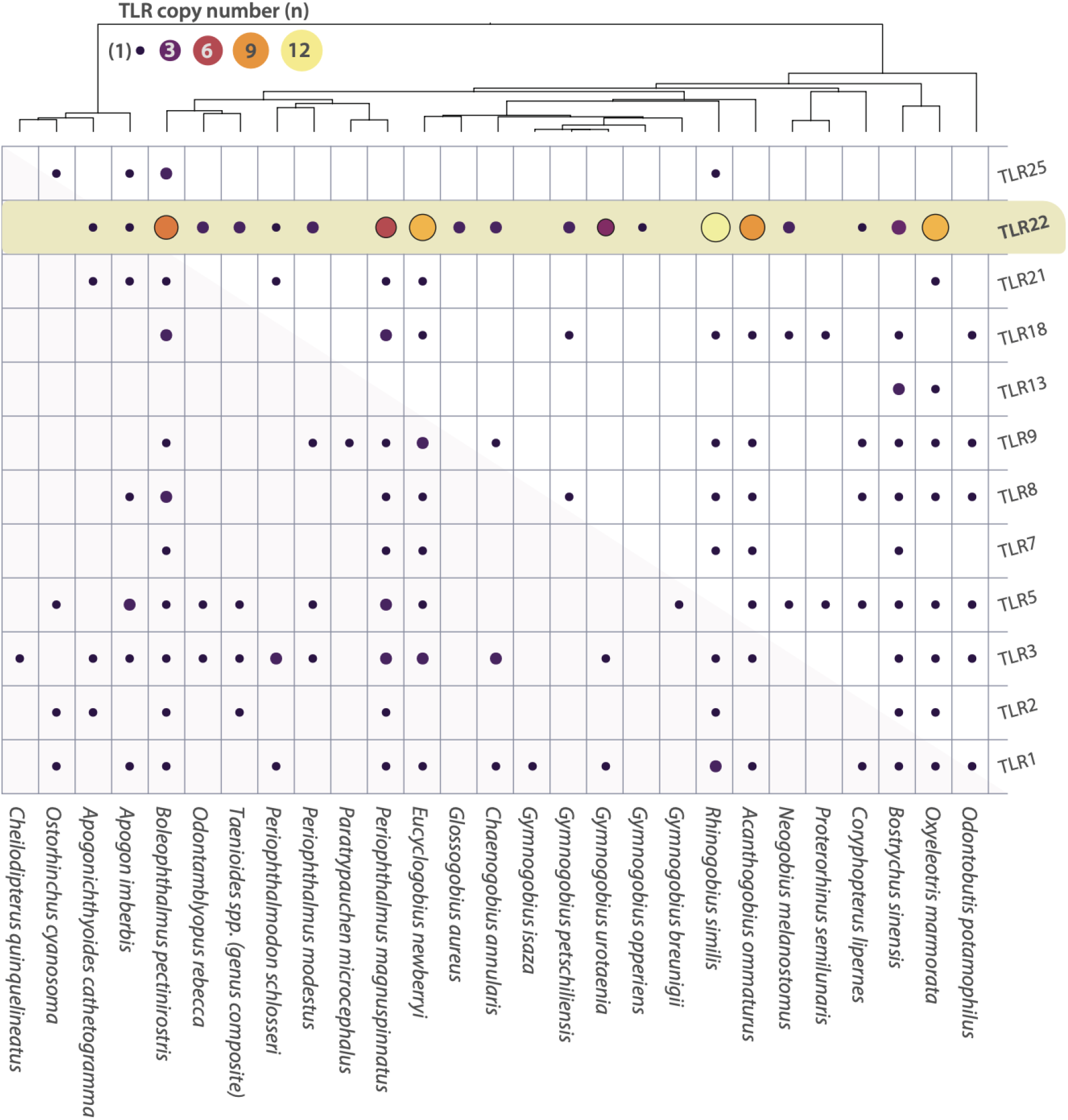
Detected variation in TLR copy number across Gobiiformes. Rows indicate TLR lineage with circle sizes and shadings corresponding to the number of unique sequences of TLR lineages between species (columns). Evolutionary relationships between species are depicted by the phylogeny at the top of the grid. Shading in the TLR22 row highlights elevated gene copy numbers relative to other TLR lineages.

**Fig. 2.**
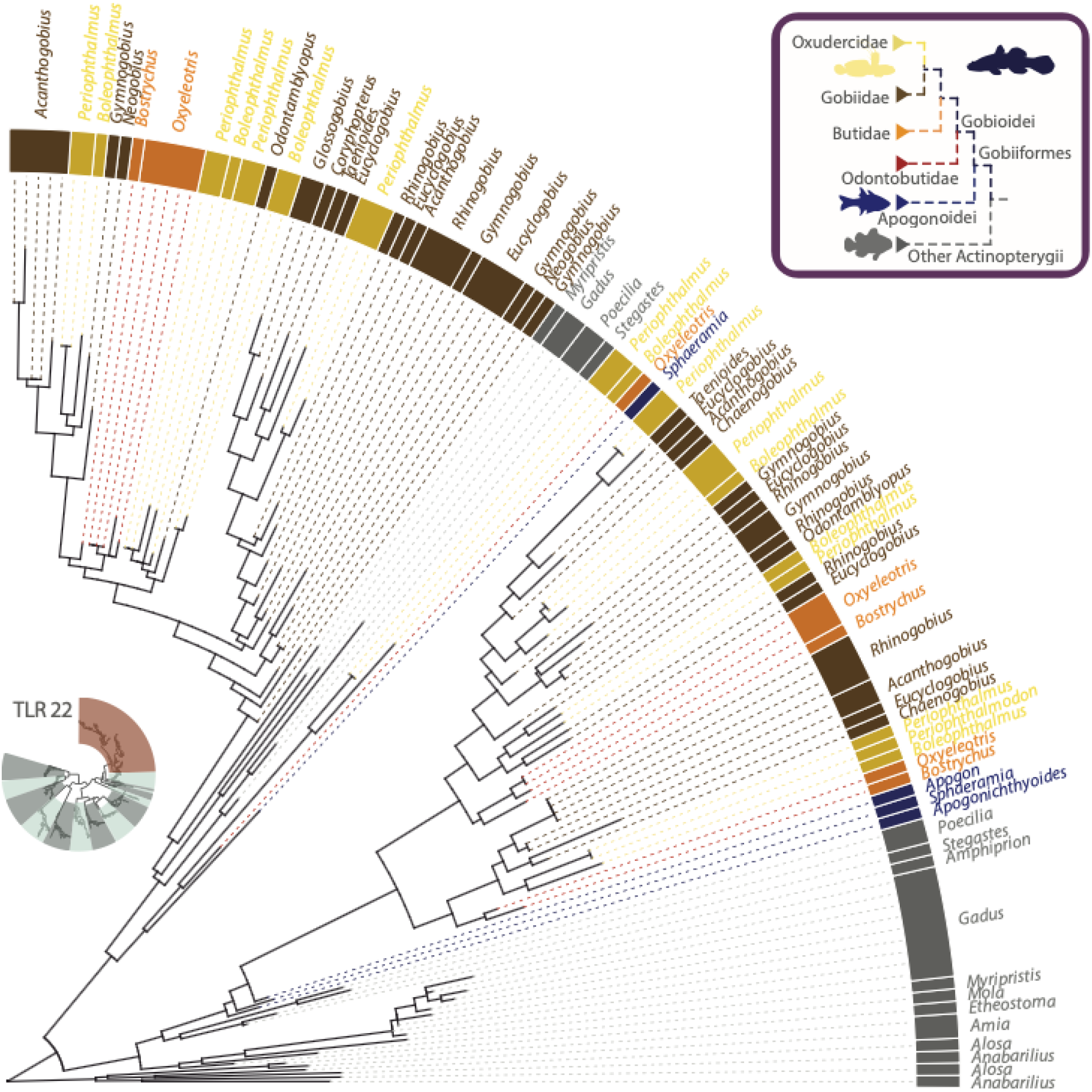
Evolutionary relationships of gobiiform TLR22 paralogs. Oxudercidae (including Mudskippers) are delimited in gold, Gobiidae by brown, Butidae by orange, Odontobutidae by red, and remaining reference actinopterygian taxa by grey. Shadings in relation to evolutionary history are included in the legend (top left). Bottom insert tree indicates the position of the TLR22 clade (warm shading) within the TLR phylogeny (**Supplemental Fig. S2**) with alternate shaded regions corresponding to other TLR subclades (**Supplemental Figs. S3-S14**). Sequence annotations are provided in **Supplemental Figs. S13a-b.**

**Fig. 3.**
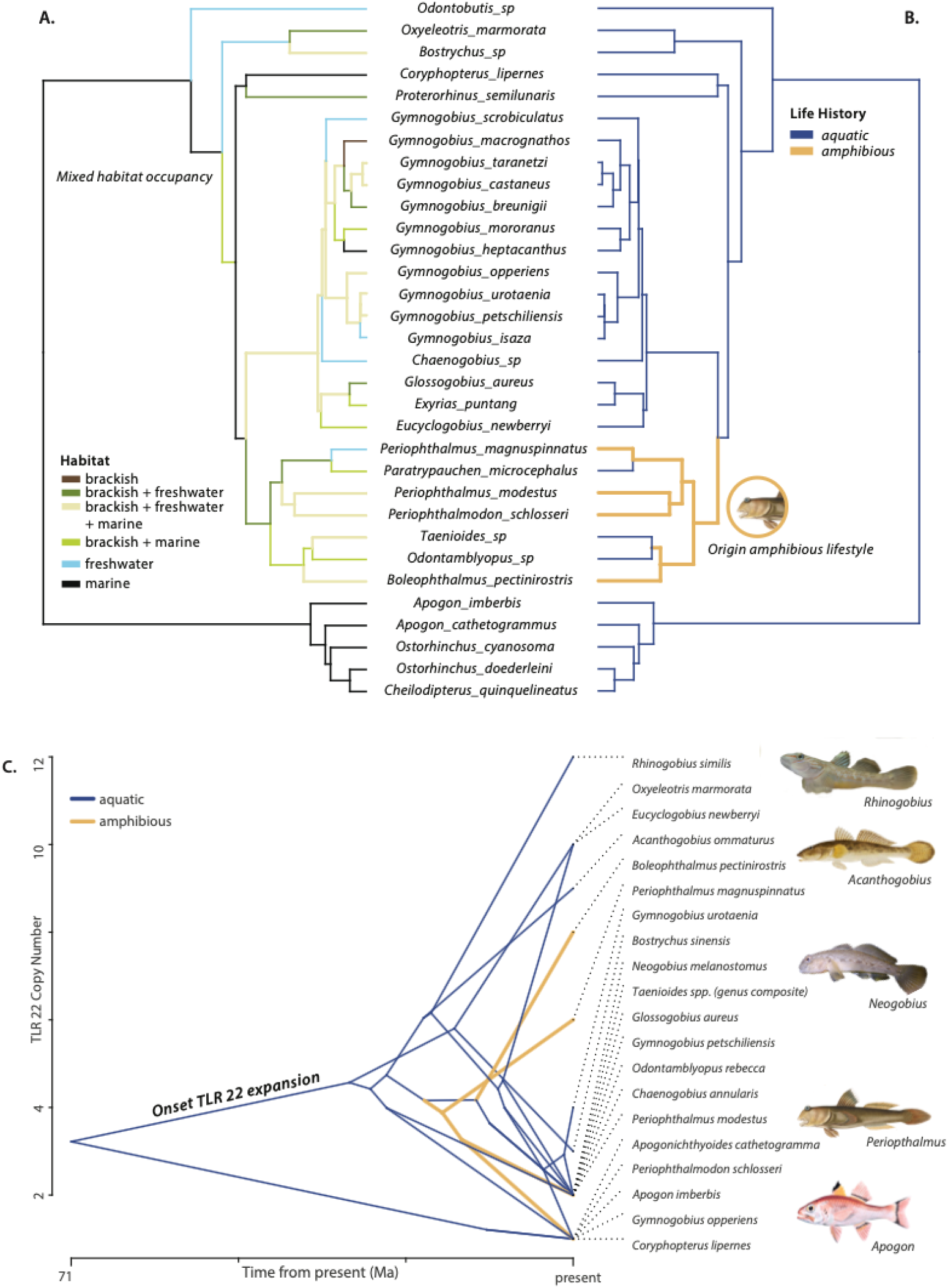
Evolutionary history of habitat transitions and TLR22 expansions. **(A)** Evolutionary history of habitat transitions across gobiiforms. Mixed habitat occupancy label indicates early evolutionary onset of gobioid fishes occupying multiple habitat types based on the ancestral state estimation. **(B)** Evolutionary transitions between amphibious and fully aquatic lifestyles. **(C).** Ancestral state estimates of TLR22 gene copies over time. Y-axis indicates TLR22 copy number, and X-axis indicates time from present day. Tips indicate extant TLR copy number variation with internal nodes representing the ancestral state estimate of TLR copy number. In all panels, shadings indicate ancestral habitat reconstruction per branch corresponding to each legend.

Within the context of pathogen defense, the expansion of TLR22 is striking. This teleost-specific receptor is consistently identified as a central component of antibacterial and antiviral immunity (Wei et al. 2023; Paria et al. 2023; Ghani et al. 2024; Su 2025), that in pufferfish is capable of signaling from the cell surface and inducing interferon-mediated antiviral responses that protect against RNA virus infection (Matsuo et al. 2008). Recent environmental virology has fundamentally revised our picture of aquatic pathogen exposure by showing that marine and freshwater systems contain a vast and still incompletely sampled virosphere. More than 4,500 novel RNA viruses were recovered from a single survey near the Yangtze River estuary in China (Wolf et al. 2020), and the finding of substantial previously unknown viral sequences has become a hallmark of recent aquatic surveys (Wolf et al. 2020; Sadeghi et al. 2021; Kolundžija et al. 2022; Urayama et al. 2024; Wu et al. 2025). However, our phylogenetic (**Fig. 2**) and ancestral state analyses (**Fig. 3**) suggest that the molecular basis of this putative antiviral receptor copy number diversity reflects an early evolutionary paralog expansion within Gobioidei (**Fig. 2**). These results suggest that gobioids have leveraged TLR22 paralogs to traverse this virosphere throughout their evolutionary history, transitioning into an exceptional range of ecological settings that span numerous marine habitats, estuaries, streams, subterranean waters, and even the terrestrial margins of aquatic ecosystems (Greenfield and Johnson 1999; Freyhof 2011; Schliewen 2011). The pathogen pressure faced by gobies across these habitats is evident even at the earliest developmental stages, where reproductive strategies such as substrate spawning or egg attachment to benthic surfaces intensify pathogen exposure and necessitate parental behaviors to mitigate larval infection such as egg fanning or cleaning, and even the secretion of antimicrobial compounds to protect embryos (Giacomello et al. 2008; Pizzolon et al. 2010; Trujillo-García et al. 2025). These early gene duplications likely acted as an immunogenetic substrate that facilitated persistence within novel pathogen-rich environments, raising the question of whether other innate immune gene families exhibit similar expansions. Moreover, this also raises the question of whether TLR repertoires in other major fish clades that occupy diverse habitats also exhibit early evolutionary paralog expansion such the one observed in Gobioidei. As more goby and other ray-finned fish genomes are sequenced, answering questions such as these will provide a powerful framework for understanding the evolutionary relationship between immunogenetic diversification and ecological breadth.

### TLR22 structural diversity is established early and remains constrained across ecological transitions

Ecological transitions are often hypothesized to promote structural innovation following gene duplication, particularly if recently duplicated paralogs undergo neofunctionalization in response to novel selective pressures. Contrary to this expectation, projection of gobiiform TLR structures into a shared protein phylomorphospace revealed that TLR22 does not occupy a unique structural region, but instead overlaps extensively with multiple other TLR subfamilies (**Fig. 4A**). This broad overlap suggests that the fundamental architecture of TLRs is highly conserved despite repeated episodes of gene duplication (**Fig. 4B**), consistent with expectations for proteins with strong structural constraints imposed by ligand recognition and downstream signaling (Goldtzvik et al. 2023). Within this conserved framework, closely related TLR22 paralogs frequently diverge in protein shape, with several lineages independently converging on similar structural solutions among lineages separated by tens of millions of years. (**Fig. 4C**). These results parallel studies of paralog evolution across diverse gene families in which localized structural changes drive sometimes dramatic functional divergences (Holub et al. 2024; Sieber et al. 2024). Accordingly, mudskippers do not occupy a distinct or expanded region relative to other taxa, a pattern supported by the absence of significant differences in morphospace occupancy between these amphibious species and other gobiiform lineages (PERMANOVA, F=0.9677, p=0.40). Together these results indicate that ecological transitions do not require the evolution of fundamentally new TLR22 architectures, but instead act upon an evolutionarily conserved structural scaffold that is repeatedly modified through localized divergence among paralogs.

**Fig. 4.**
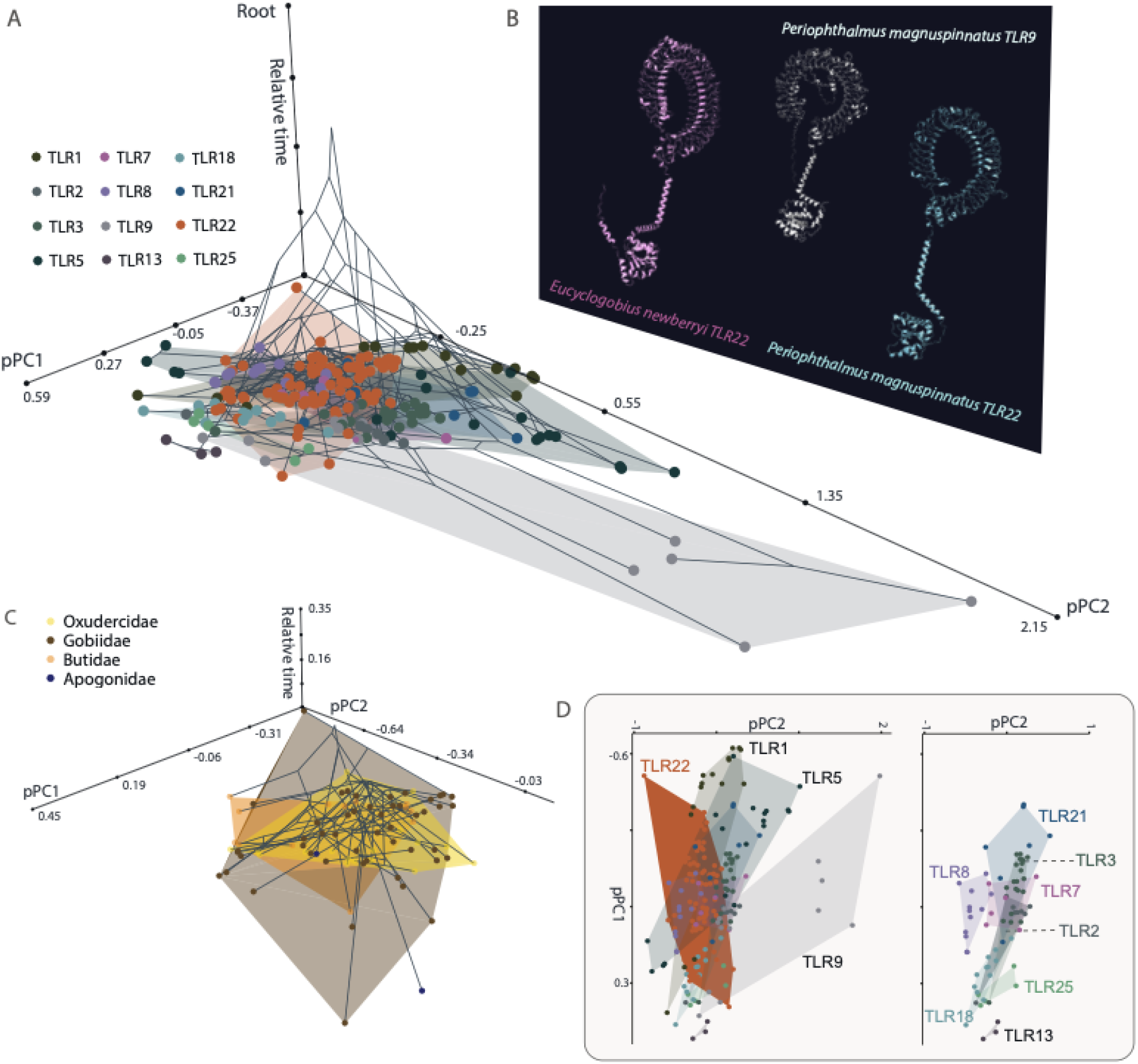
Projection of evolutionary relationships into protein phylomorphospace. **(A)** Phylomorphospace of the principal components axes of the full-length TLR shapes predicted by ESMfold. The X and Y axes represent PC1 (68.3 %variance explained) and PC2 (19.9 % variance explained; see **Supplemental Fig. S16**) of the protein shape analysis, with circles indicating proteins colored by TLR clade. Connecting lines on the Z axis reflect the evolutionary history of the TLR paralogs scaled to a relative height of 1.0. Internal nodes represent the ancestral shape estimate. Shaded polygons indicate the convex hull of each respective TLR clade. **(B)** Example protein structures predicted by ESMfold. **(C)** The TLR22 phylomorphospace isolated from panel A, colored by taxonomic family. **(D)** The X and Y axis from panel A, with specific TLRs isolated to interpret visual interpretation and delimitation of morphospace convex hulls.

If immune gene paralogs followed a simple adaptive radiation model, it may be expected that immune gene families depict a signature of rapid receptor structure changes as lineages encounter novel pathogen regimes. However, the mode of TLR22 evolution we observe is instead characterized by the retention of core receptor architectures (**Figs. 4A** and **4D**) and subsequent diversification through fine-scale modification within those inherited regimes. This interpretation is strongly supported by the molecular immunology of TLR22, where modest changes can produce disproportionately large functional consequences. In grass carp, duplicated TLR22 paralogs (CiTLR22a and CiTLR22b) exhibit antagonistic functional effects. CiTLR22a enhances antiviral signaling through the activation of interferon and NFκB pathways whereas CiTLR22b suppresses both pathways and permits viral propagation (Ji et al. 2019). This antagonistic functional difference is striking when considering the shared core architecture of these receptors including localization to acidic endolysosomal compartments, binding dsRNA analogs only under low pH conditions, and signaling through MyD88-dependent pathways (Ji et al. 2019). Fine-scale immunogenetic tuning is further supported by comparative work in cyprinids, where barbel chub and grass carp TLR22 differ only modestly in sequence, yet substitutions concentrated in the LRR ectodomain and a small number of TIR-associated residues alter IFN-β induction and viral resistance, and domain-swapping experiments confirm that these localized changes are sufficient to shift antiviral performance (Wang et al. 2017). These patterns are mechanistically consistent with the modular architecture of TLRs, in which the LRR domains define ligand interaction surfaces while the TIR domain preserves downstream signaling capacity, allowing ligand recognition to evolve through incremental modification of receptor surfaces rather than wholesale structural innovation. These results indicate that ecological transitions do not require major structural shifts of TLR22, but instead draw on ancient receptor architectures that are progressively refined to generate functional diversity among closely related paralogs.

### TLR diversification proceeds through modular reconfiguration of conserved receptor architecture

The extracellular and cytoplasmic domains of immune receptors are expected to experience distinct evolutionary pressures because they, respectively, often mediate ligand recognition and downstream signaling (Guida et al. 1992; Filip and Mundy 2004; Mikami et al. 2012; Hilbert et al. 2023). However, our analyses suggest that variation in LRR organization within the extracellular region is not randomly distributed among TLR22 paralogs (**Fig. 5**). Instead, LRRs occupy broadly conserved, interspersed positions across receptor sequences. This pattern is consistent with the canonical organization of LRR-containing proteins, in which conserved leucine residues stabilize the β-sheet/α-helix scaffold while solvent-exposed positions within the repeat units form the ligand-binding surface (Bengtsson et al. 1995; Jang et al. 2021). The conservation of LRR positions across paralogs suggests that these repeat domains provide a modular framework in which receptor architecture can diversify without disrupting the broader receptor organization, a pattern also observed in other LRR-containing proteins (Metcalfe et al. 2023). In this case, diversification of LRR arrays is expected to modify interactions with pathogen-associated molecular patterns while preserving the conserved structural scaffold required for receptor function.

**Fig. 5.**
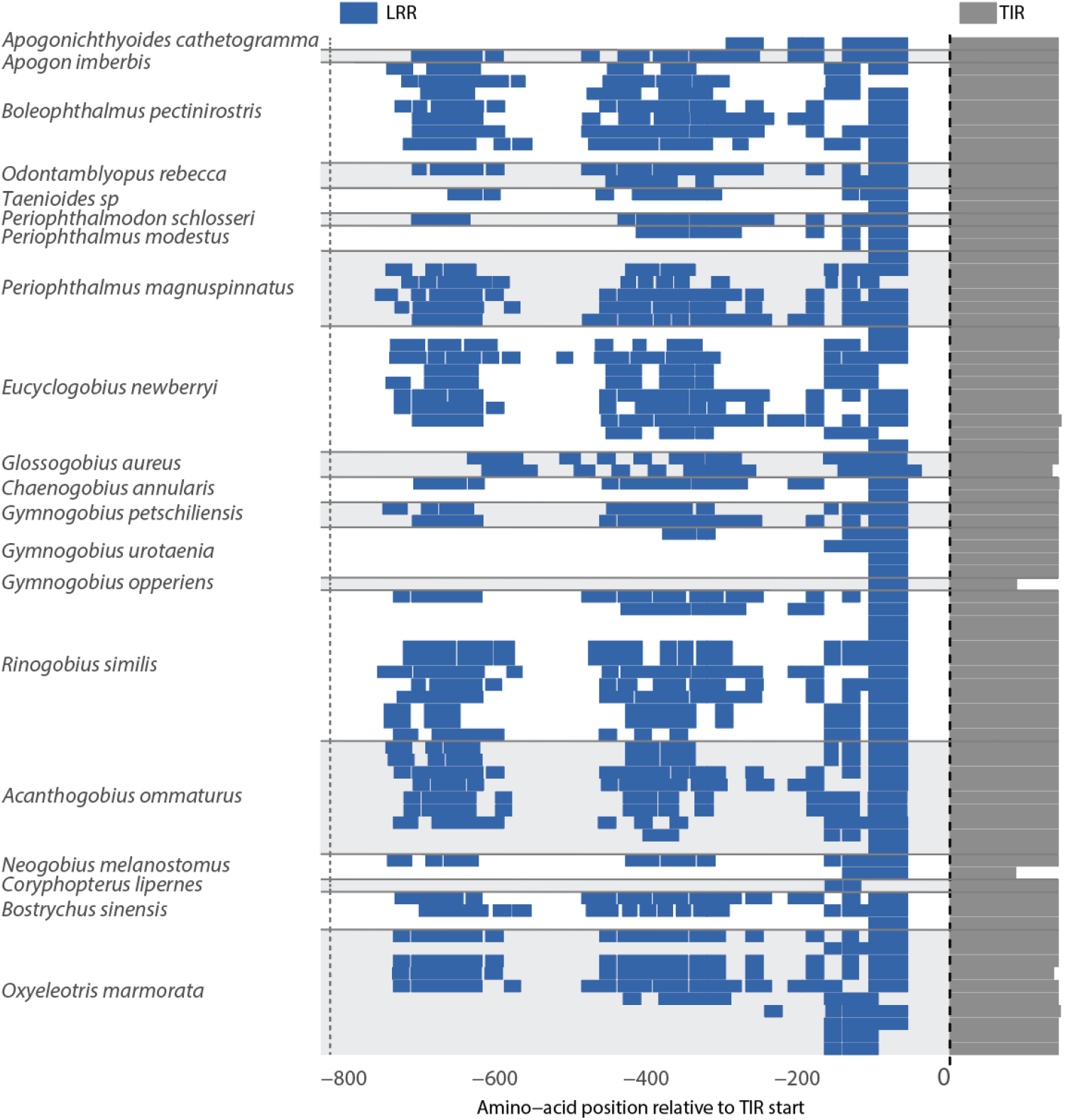
Variation of TLR22 LRR domain architecture and sequence composition. Map of the TLR22 LRR and TIR domain positions, centered at the start of the TIR domain. Rows correspond to individual TLR22 genes with shadings indicating individual LRR domains or TIR domains organized by species in which they occur.

We found variation in ectodomain size was closely coupled to expansion and contraction of the LRR array. Across TLRs, extracellular-region length increased with the number of annotated LRRs (**Fig. 6A**), a relationship strongly supported by phylogenetic generalized least-squares regression (P < 0.0001; **Fig. 6B**). This association remained strongly supported after accounting for total protein length, with residual extracellular-region length positively associated with residual LRR number (P < 0.0001 ; **Fig. 6C**). Residual LRR number was also tightly correlated with the residual combined length of all LRRs (P < 0.0001; **Fig. 6D**), confirming that increases in ectodomain size primarily reflect the addition or enlargement of recognizable repeat units. In contrast, residual non-LRR extracellular length was negatively associated with residual LRR number after accounting for total protein length (P < 0.0001; **Fig. 6E**). Thus, expansion of the LRR array does not appear to require proportional expansion of the intervening extracellular sequence. Instead, TLR evolution proceeds through the insertion, deletion, or modification of repeat domains, providing a structural mechanism through which ligand recognition can diversify without disrupting the conserved receptor architecture.

**Fig. 6.**
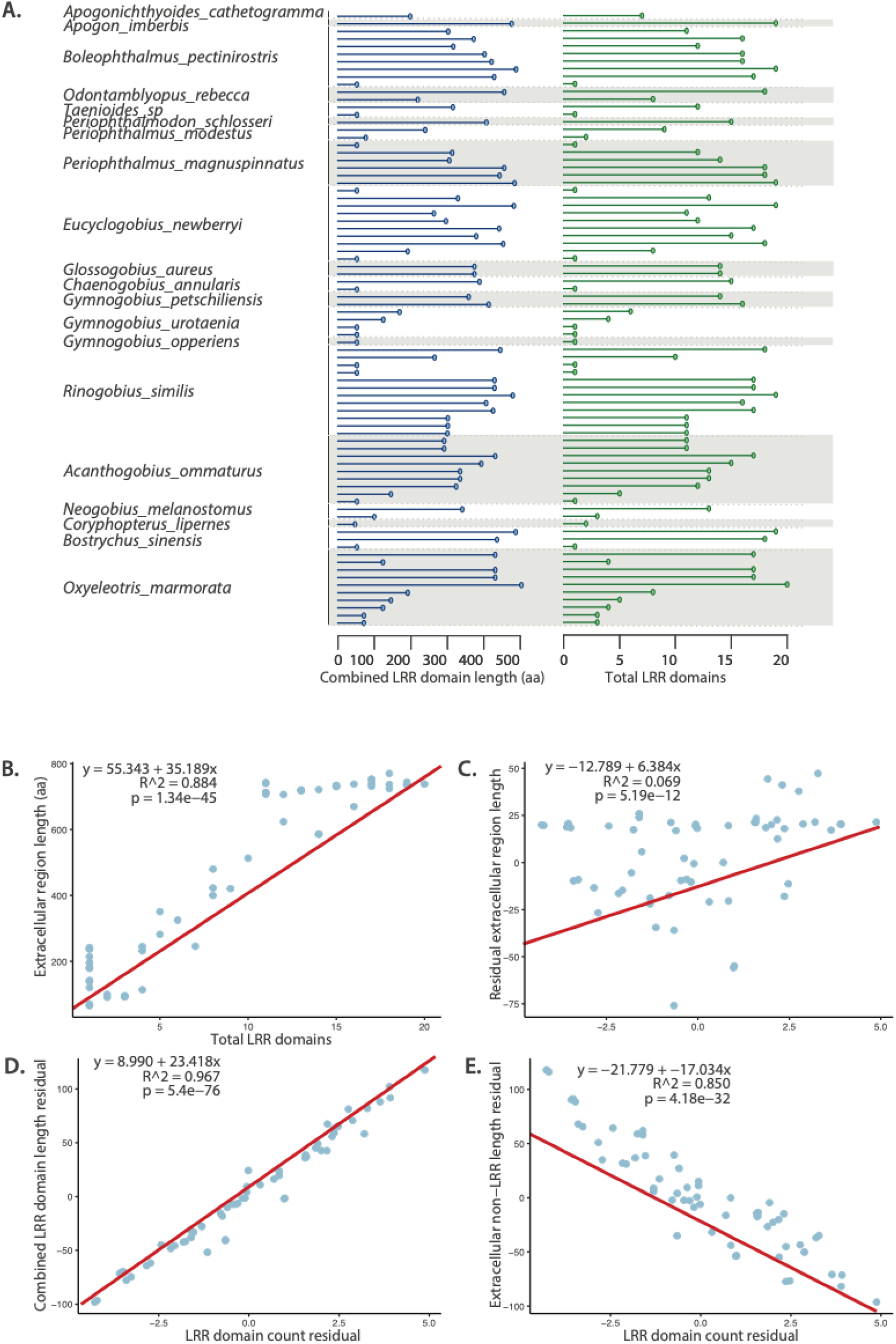
Evolutionary variability of TLR22 LRR repeats. (**A)** Lollipop plots of the combined length of the LRR domains within each sequence (left) compared with the number of LRR domains (right). Horizontal dotted lines delineate sequences that occur with labelled species (**B)** The relationship between the size of the extracellular region and the number of LRR domains. (**C)** The relationship between the residual length of the extracellular region and the residual number of LRR domains when accounting for total sequence length. (**D)** The relationship between the residual total length of the LRR domains combined and the residual number of LRR domains when accounting for total sequence length. (**E)** The relationship between the residual length of the extracellular region excluding LRR domains and the residual number of LRR domains when accounting for total sequence length. Red line indicates best-fit line from a phylogenetic least squares regression with the regression equation, R-squared value, and p value indicated in each panel.

Across all TLRs, LRR length exhibits strong phylogenetic signal, with Pagel’s λ near 1 (λ = 0.96, p = 7.34 × 10^−42^), indicating that variation in LRR architecture is highly structured by shared evolutionary history. This signal sharply departs from Brownian expectations, as Blomberg’s K is effectively zero (K = 5.2 × 10⁻⁶, p = 0.235). The combination of high λ and negligible K suggests that LRR length evolution is phylogenetically constrained but does not follow a gradual process. Instead, changes in repeat architecture may be concentrated along particular branches and subsequently retained or repeatedly re-evolved within descendant lineages. Consistent with this interpretation, ancestral-state reconstructions revealed that members of each major TLR clade generally varied within a characteristic range of LRR numbers rather than exploring the complete range observed across the family (**Fig. 7**).

**Fig. 7.**
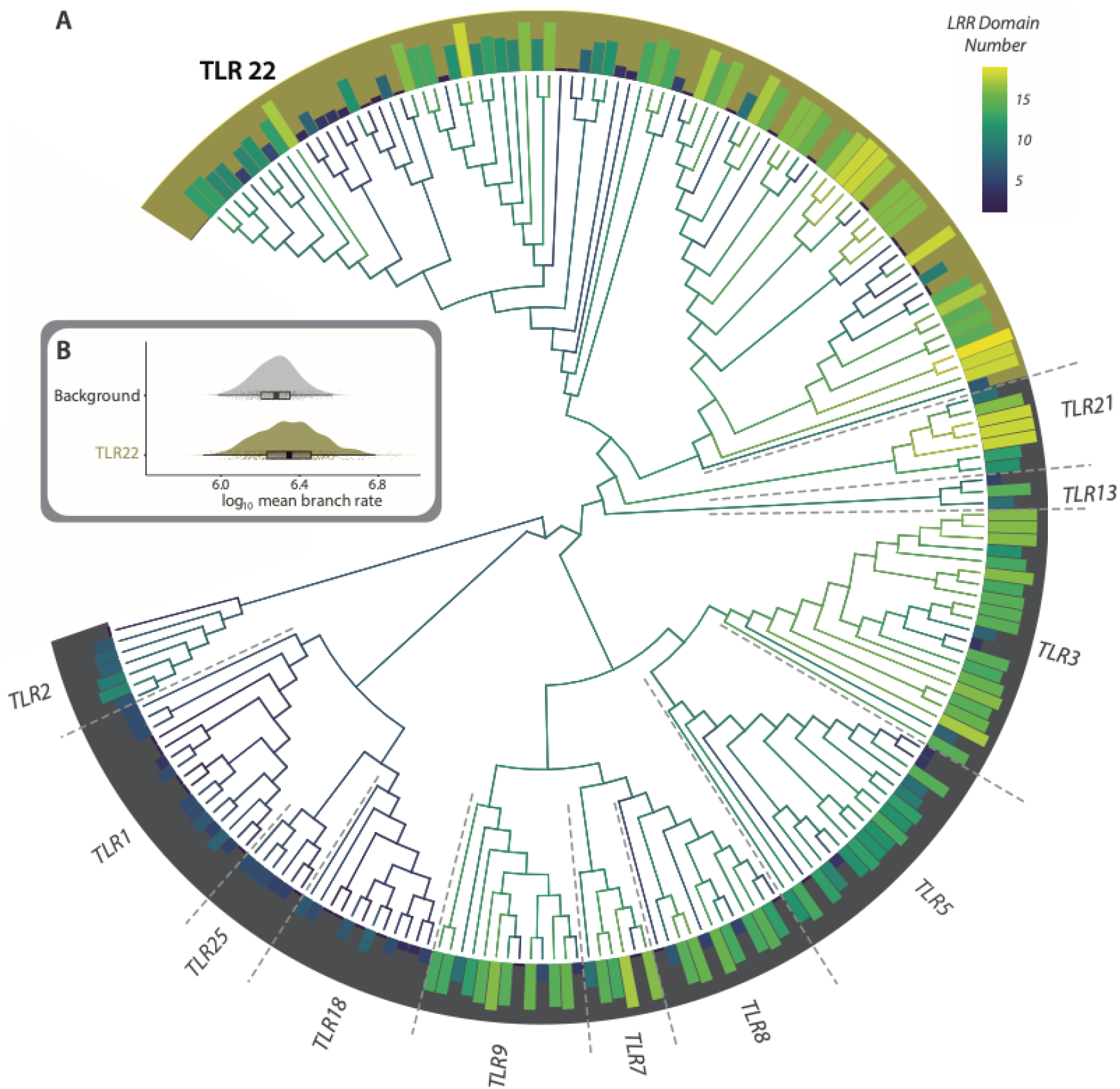
Evolutionary variability of TLR22 LRR repeats. **(A)** Ancestral State estimations of LRR domain copy number (**Supplemental Datasheet 1**) across the phylogeny of TIR domains of TLR paralogs from Gobiiformes. The number of extant and estimated ancestral LRR domains per paralog correspond to the shaded histograms and branch lengths, respectively. TLR clades are delineated by the outer band with TLR22 highlighted in gold. **(B)** 95% Highest posterior density interval of the log-transformed mean diversification rate per branch between TLR22 and all other TLRs (background).

TLR22 departed markedly from this family-wide pattern. Neither Pagel’s λ (λ = 7.33 × 10⁻⁵, *P* = 0.99) nor Blomberg’s *K* (*K* = 2.28 × 10⁻⁵, *P* = 0.43) supported detectable phylogenetic signal in LRR architecture, indicating that closely related paralogs do not consistently retain similar repeat organizations. However, shifts in LRR architecture were not associated with rapid evolutionary lability. Instead, our BAMM analyses indicated that the average rate of LRR expansion and contraction within TLR22 was indistinguishable from the background rate estimated across TLRs (**Fig. 7**). This suggests that the architectural diversity of TLR22 did not arise from accelerated structural evolution. Instead, it appears to reflect repeated reconfiguration of LRR architecture under the same evolutionary processes operating throughout all TLRs. Gene duplication therefore increases the number of receptor copies available for selection rather than increasing the rate at which structural variation is generated (Kuzmin et al. 2022; Deem et al. 2024). Collectively, these results suggest that ecological transitions need not be accompanied by accelerated structural innovation. Rather, expanded paralog repertoires permit repeated exploration of existing architectural possibilities while preserving the broader evolutionary dynamics governing TLR diversification.

### Conclusion

Evolutionary transitions are often interpreted through the lens of contemporaneous innovation, with trait diversification events viewed as responses to newly encountered ecological pressures (Dornburg et al. 2011; Parker et al. 2022; Porto et al. 2025). However, our analyses of gobiiform TLRs indicate that much of the structural and potential functional diversity relevant to pathogen recognition was established prior to many of the ecological transitions that define this radiation. Across phylogenetic, structural, and sequence-level analyses, we consistently recover a pattern in which early paralog expansions partitioned receptor diversity into discrete structural regimes that have persisted over tens of millions of years. Subsequent evolution primarily proceeded through localized reconfiguration of ligand-recognition architecture rather than repeated invention of fundamentally new receptor forms. Collectively, these results resolve the temporal ambiguity surrounding immune gene diversification in this clade, supporting a history in which ancient immunogenetic diversity preceded, and may have subsequently facilitated ecological transitions rather than arising as a consequence of them. More broadly, our findings suggest that the adaptive consequences of gene family diversification may unfold long after duplication events themselves, as ancient receptor diversity is repeatedly refined and redeployed as lineages encounter new ecological challenges.

## Acknowledgements

We thank members of the Dornburg and Yoder labs for helpful comments during the early stages of this project.

## Data availability

Custom scripts, protein structures, alignments, raw sequences, and phylogenetic outputs are available on Zenodo (DOI: 10.5281/zenodo.21628786; https://zenodo.org/records/21628787)

## Supplemental Materials

### Overview

In addition to this supplemental document, this manuscript includes two supplemental datasheets to enhance reproducibility. **Supplemental Datasheet 1** contains all TLR sequences, TIR domains, LRR domains, and Accession Numbers. **Supplemental Datasheet 2** is an interactive html document that provides a graphical representation of TLRs used in this study and confident predictions of domain architecture.

### Supplemental Figures

**Supplemental Fig. S1.**
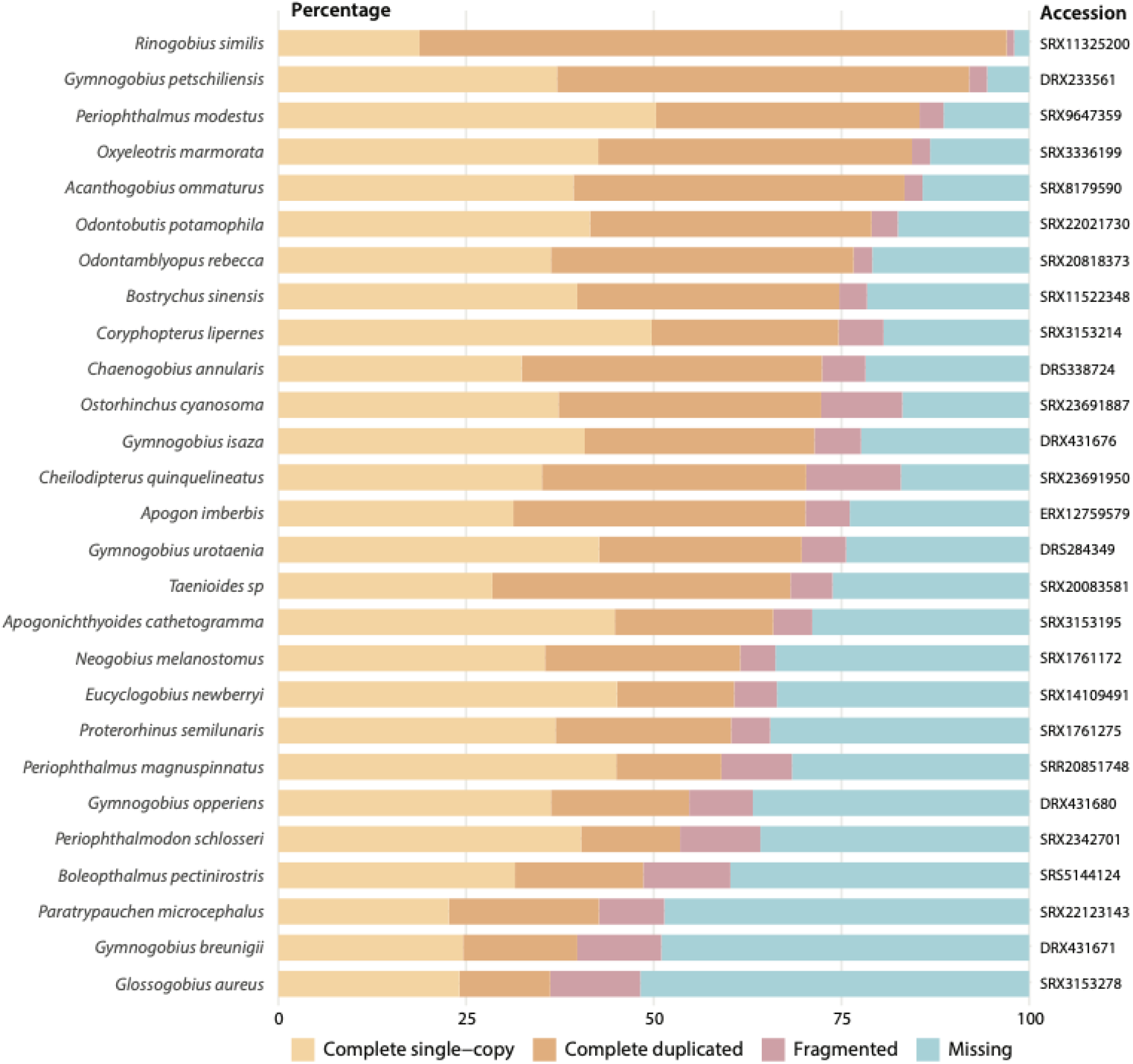
BUSCO scores of transcriptomes assembled for analyses. Shading corresponds to percent completeness with cooler colors indicating complete single copy (cream), duplicate (peach), fragmented (purple), or missing (blue) loci. Accession refers to the Sequence Read Archive (SRA) reference number on GenBank.

**Supplemental Fig. S2.**
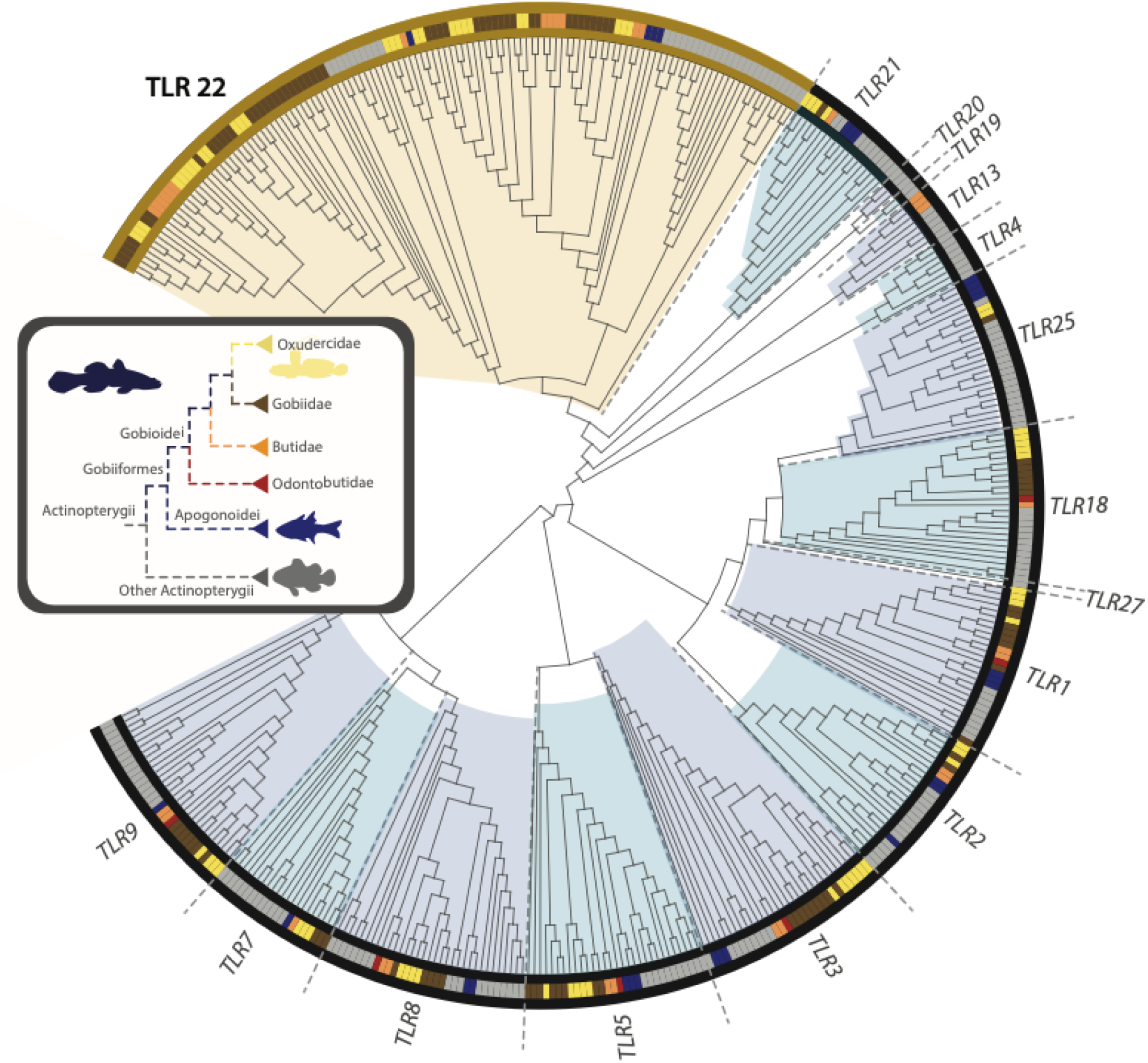
TLR paralogs across the Actinopterygiian phylogeny. Phylogeny of TIR domains of TLR paralogs from Gobiiformes and reference Actinopterygian taxa (**Supplemental Datasheet 1**) estimated using Maximum likelihood. Oxudercidae (including Mudskippers) are delimited in gold, Gobiidae by brown, Butidae by orange, Odontobutidae by red, and remaining reference actinopterygian taxa by grey. Bootstrap support values 70 or higher are indicated at nodes as indicated in the legend. A detailed view of the TLR22 clade is provided in Fig. 2. Alternate shaded regions correspond to other TLR subclades.

**Supplemental Fig. S3.**
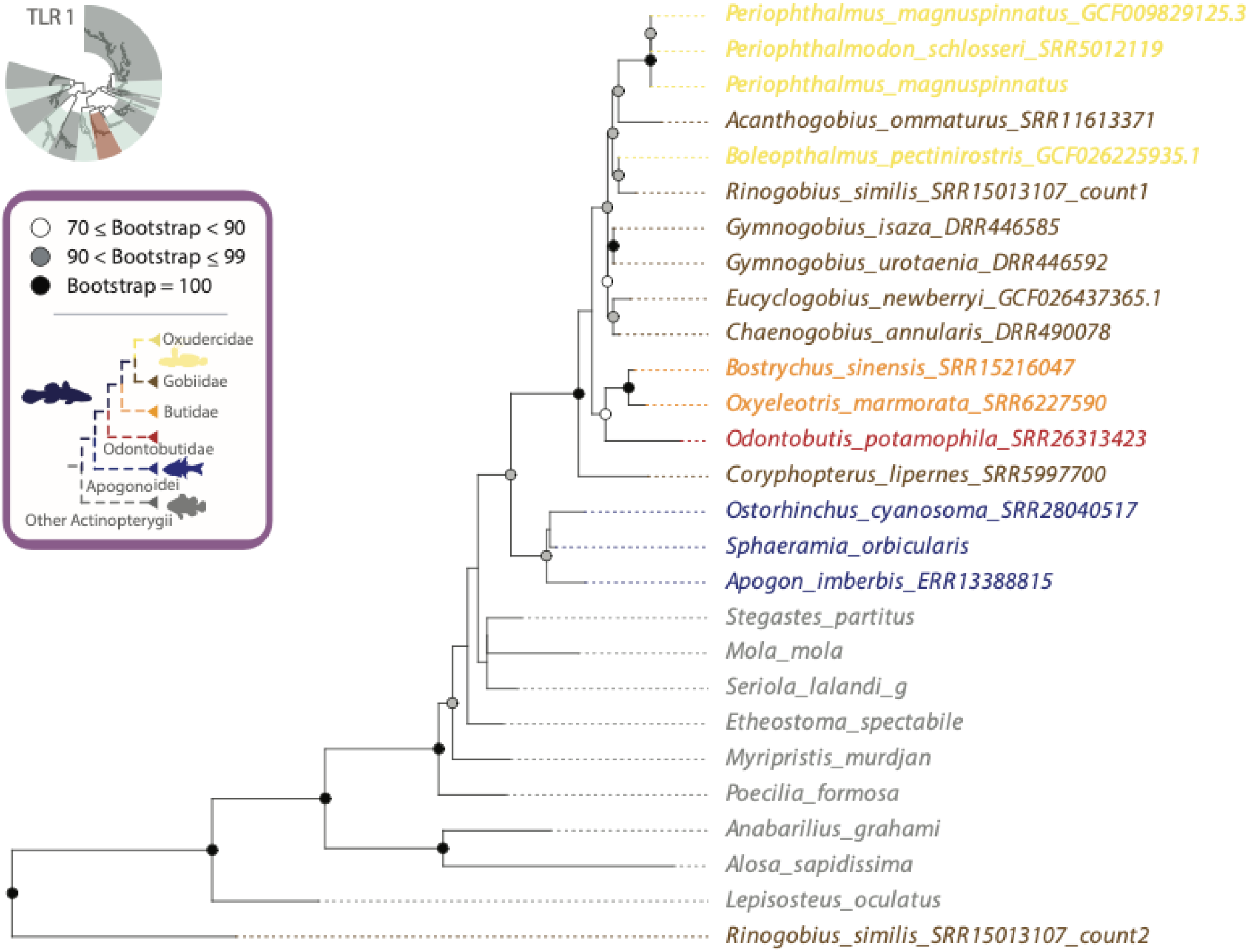
Evolutionary relationships of Gobiiform TLR1 paralogs. Detailed view of the TLR1 phylogeny in **Supplemental Fig. S2**. Oxudercidae (including Mudskippers) are delimited in gold, Gobiidae by brown, Butidae by orange, Odontobutidae by red, and remaining reference actinopterygian taxa by grey. Bootstrap support values 70 or higher are indicated at nodes as indicated in the legend. Sequences without accession numbers were obtained from the TIR dataset (using suggested nomenclature) from Supplemental Table S1 of Carlson et al. (2023). Single letter suffixes in taxon names were used to differentiate taxon names and correspond to the tree file in the data archive.

**Supplemental Fig. S4.**
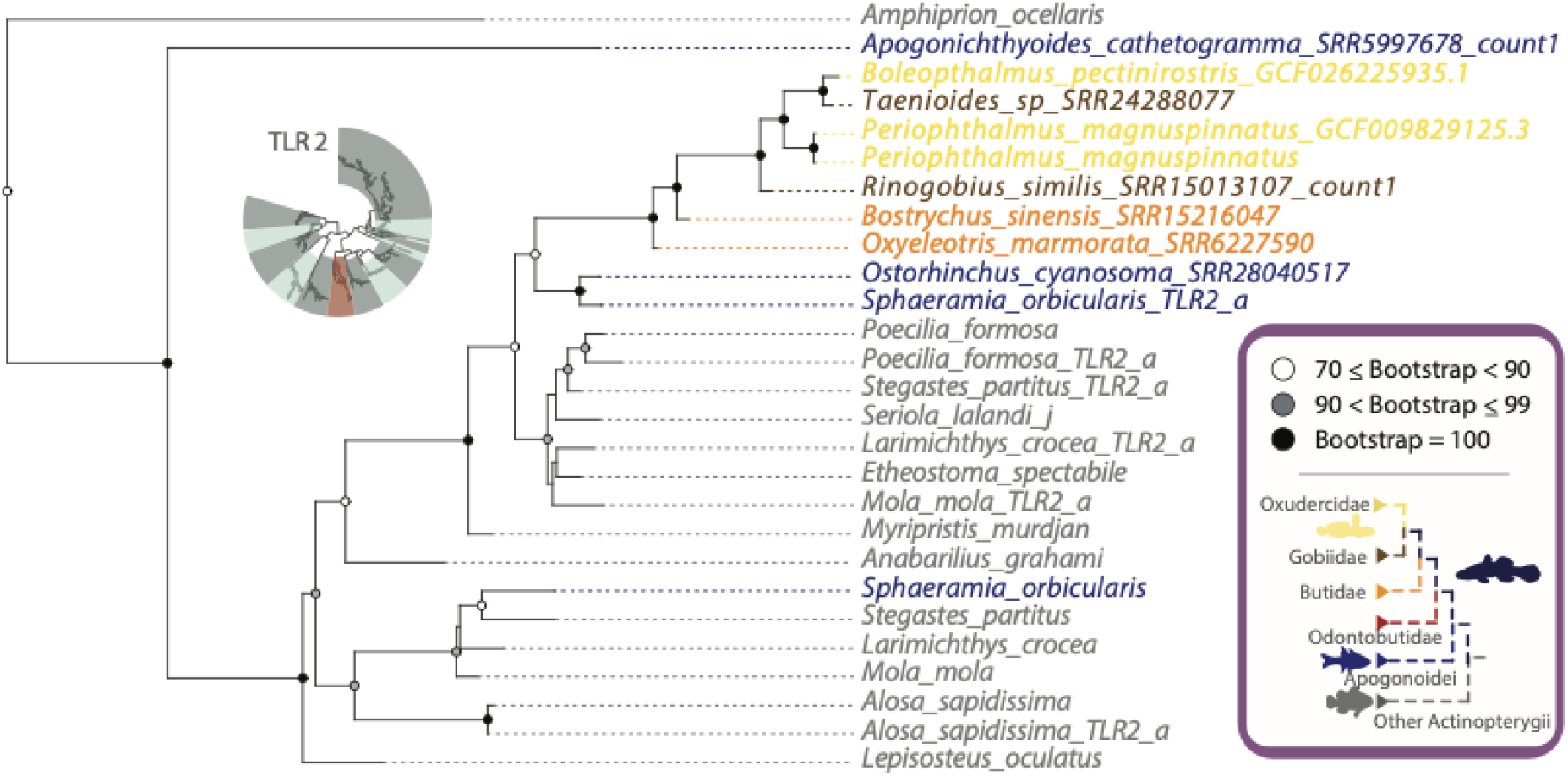
Evolutionary relationships of Gobiiform TLR2 paralogs. Detailed view of the TLR2 phylogeny in **Supplemental Fig. S2**. Oxudercidae (including Mudskippers) are delimited in gold, Gobiidae by brown, Butidae by orange, Odontobutidae by red, and remaining reference actinopterygian taxa by grey. Bootstrap support values 70 or higher are indicated at nodes as indicated in the legend. Sequences without accession numbers were obtained from the TIR dataset (using suggested nomenclature) from Supplemental Table S1 of Carlson et al. (2023). Single letter suffixes in taxon names were used to differentiate taxon names and correspond to the tree file in the data archive.

**Supplemental Fig. S5.**
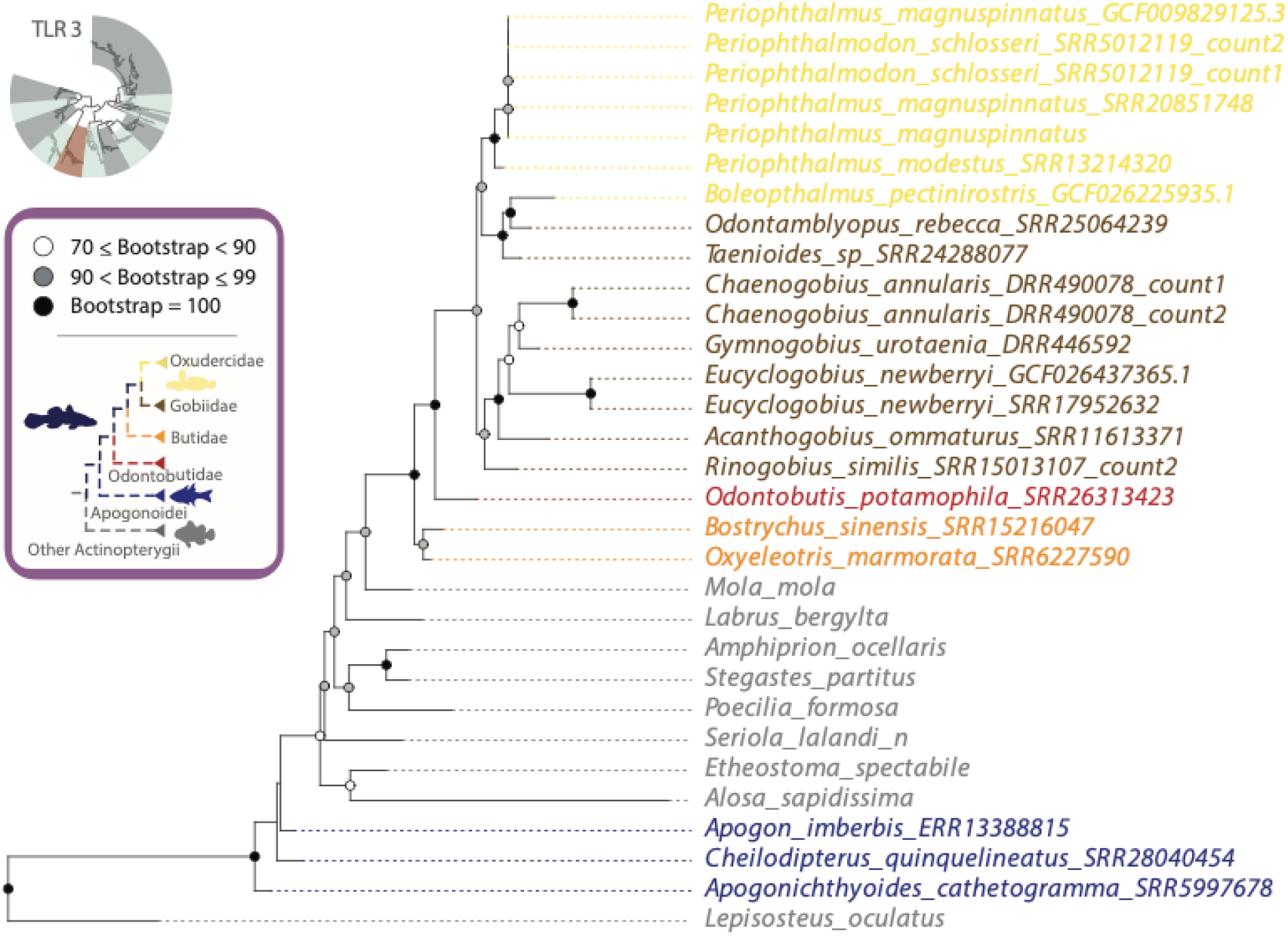
Evolutionary relationships of Gobiiform TLR3 paralogs. Detailed view of the TLR3 phylogeny in **Supplemental Fig. S2**. Oxudercidae (including Mudskippers) are delimited in gold, Gobiidae by brown, Butidae by orange, Odontobutidae by red, and remaining reference actinopterygian taxa by grey.Bootstrap support values 70 or higher are indicated at nodes as indicated in the legend. Sequences without accession numbers were obtained from the TIR dataset (using suggested nomenclature) from Supplemental Table S1 of Carlson et al. (2023). Single letter suffixes in taxon names were used to differentiate taxon names and correspond to the tree file in the data archive.

**Supplemental Fig. S6.**
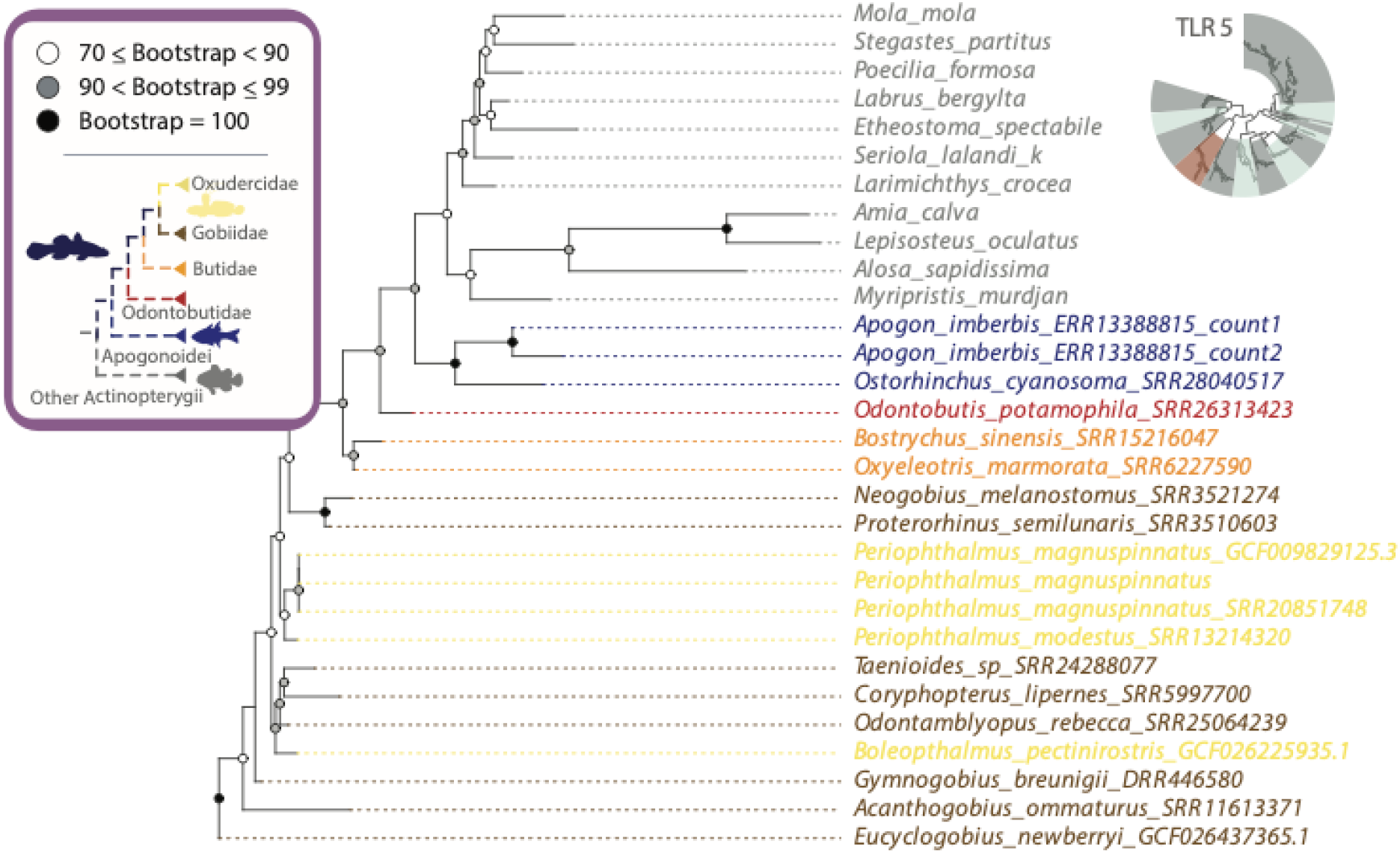
Evolutionary relationships of Gobiiform TLR5 paralogs. Detailed view of the TLR5 phylogeny in **Supplemental Fig. S2**. Oxudercidae (including Mudskippers) are delimited in gold, Gobiidae by brown, Butidae by orange, Odontobutidae by red, and remaining reference actinopterygian taxa by grey. Bootstrap support values 70 or higher are indicated at nodes as indicated in the legend. Sequences without accession numbers were obtained from the TIR dataset (using suggested nomenclature) from Supplemental Table S1 of Carlson et al. (2023). Single letter suffixes in taxon names were used to differentiate taxon names and correspond to the tree file in the data archive.

**Supplemental Fig. S7.**
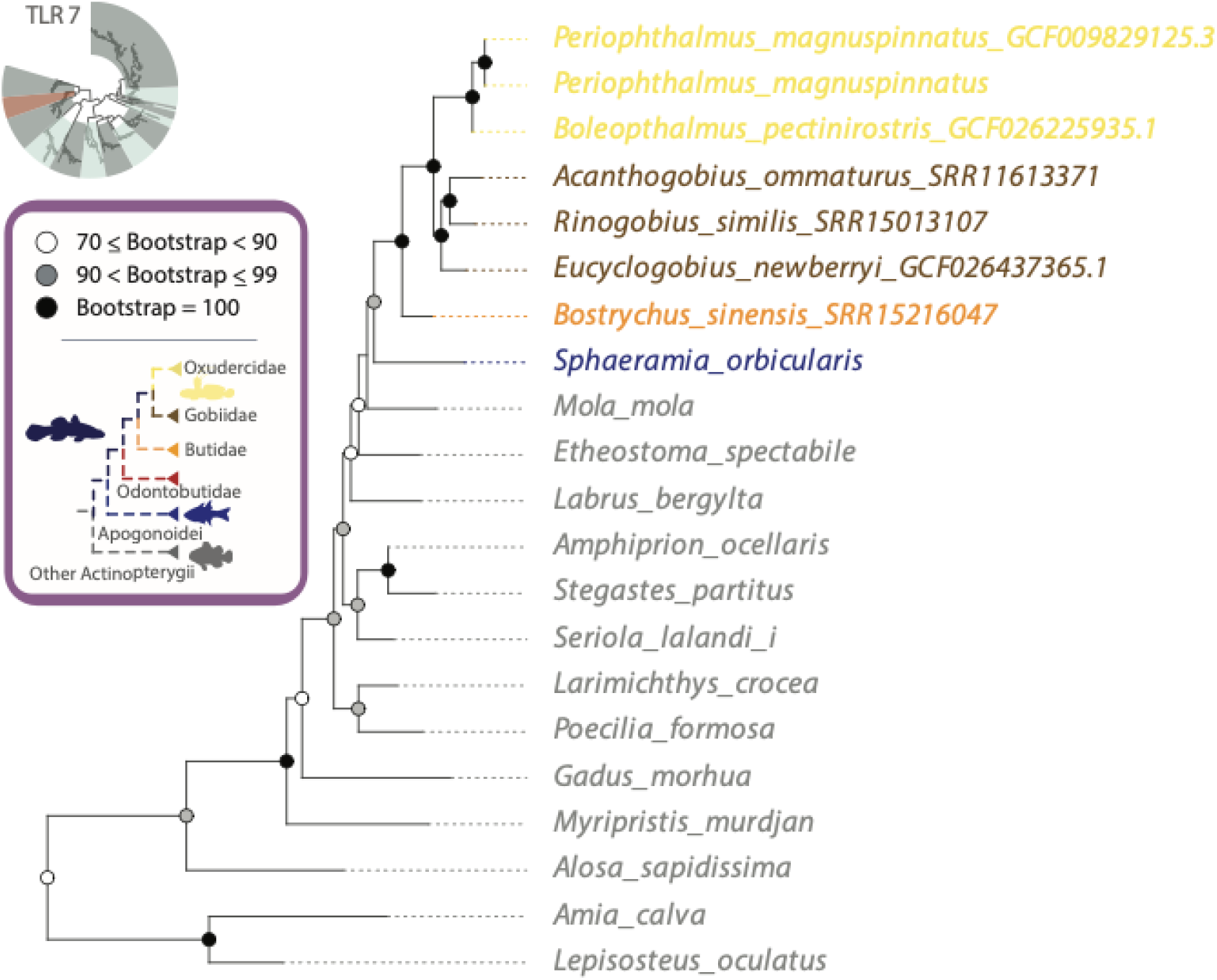
Evolutionary relationships of Gobiiform TLR7 paralogs. Detailed view of the TLR7 phylogeny in **Supplemental Fig. S2**. Oxudercidae (including Mudskippers) are delimited in gold, Gobiidae by brown, Butidae by orange, Odontobutidae by red, and remaining reference actinopterygian taxa by grey. Bootstrap support values 70 or higher are indicated at nodes as indicated in the legend. Bootstrap support values 70 or higher are indicated at nodes as indicated in the legend. Sequences without accession numbers were obtained from the TIR dataset (using suggested nomenclature) from Supplemental Table S1 of Carlson et al. (2023). Single letter suffixes in taxon names were used to differentiate taxon names and correspond to the tree file in the data archive.

**Supplemental Fig. S8.**
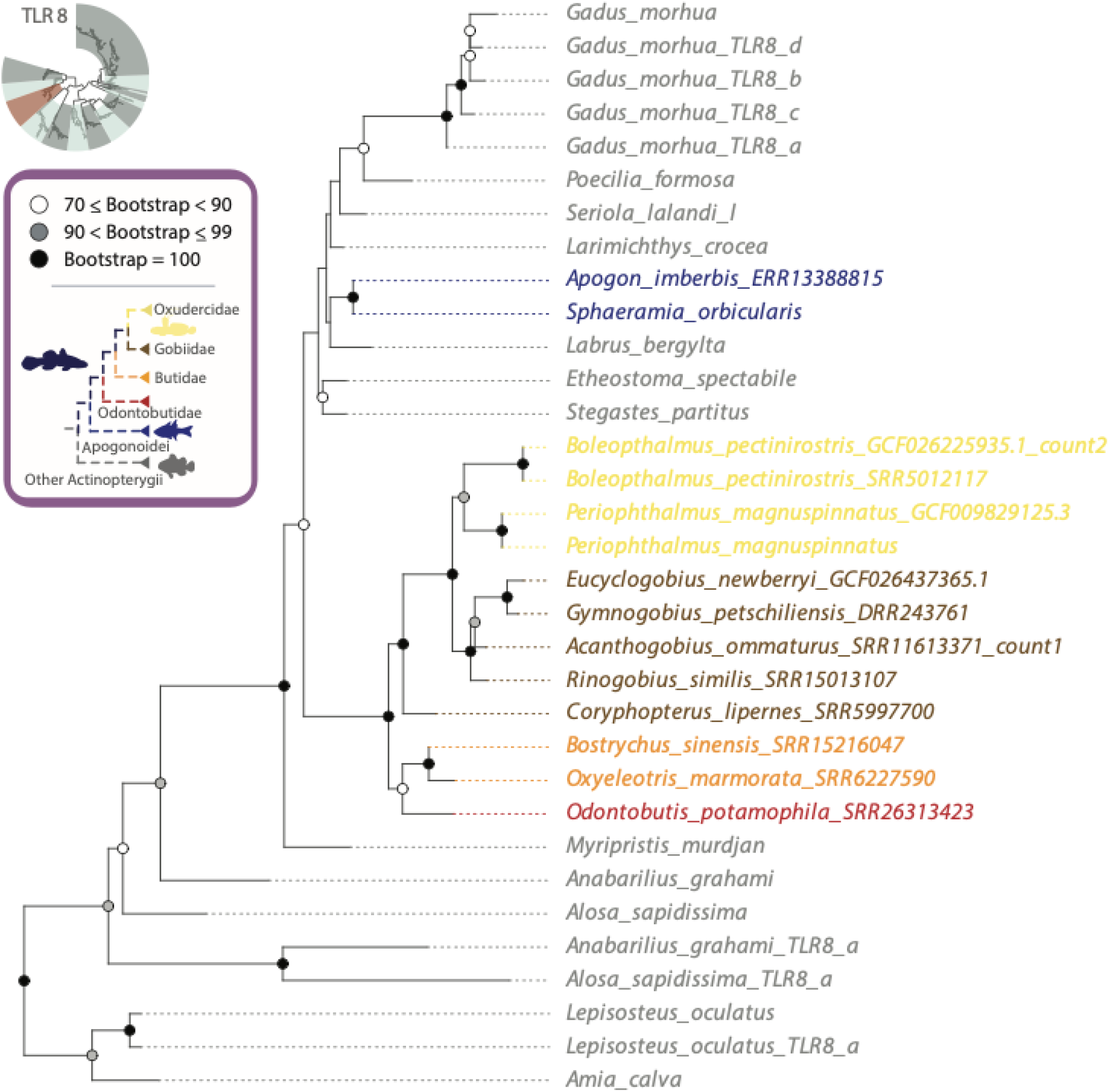
Evolutionary relationships of Gobiiform TLR8 paralogs. Detailed view of the TLR8 phylogeny in **Supplemental Fig. S2**. Oxudercidae (including Mudskippers) are delimited in gold, Gobiidae by brown, Butidae by orange, Odontobutidae by red, and remaining reference actinopterygian taxa by grey. Bootstrap support values 70 or higher are indicated at nodes as indicated in the legend. Sequences without accession numbers were obtained from the TIR dataset (using suggested nomenclature) from Supplemental Table S1 of Carlson et al. (2023). Single letter suffixes in taxon names were used to differentiate taxon names and correspond to the tree file in the data archive.

**Supplemental Fig. S9.**
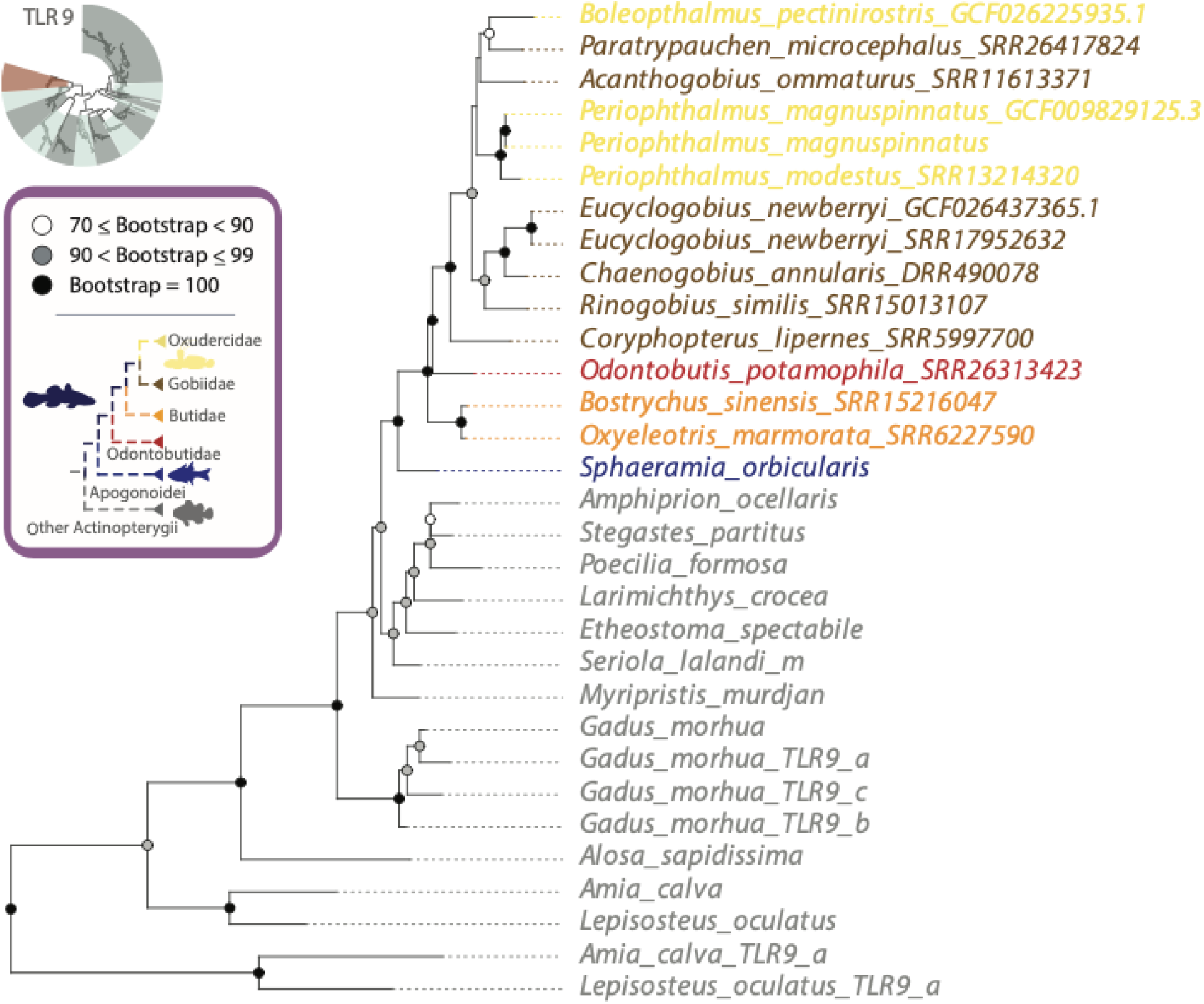
Evolutionary relationships of Gobiiform TLR9 paralogs. Detailed view of the TLR9 phylogeny in **Supplemental Fig. S2**. Oxudercidae (including Mudskippers) are delimited in gold, Gobiidae by brown, Butidae by orange, Odontobutidae by red, and remaining reference actinopterygian taxa by grey. Bootstrap support values 70 or higher are indicated at nodes as indicated in the legend. Sequences without accession numbers were obtained from the TIR dataset (using suggested nomenclature) from Supplemental Table S1 of Carlson et al. (2023). Single letter suffixes in taxon names were used to differentiate taxon names and correspond to the tree file in the data archive.

**Supplemental Fig. S10.**
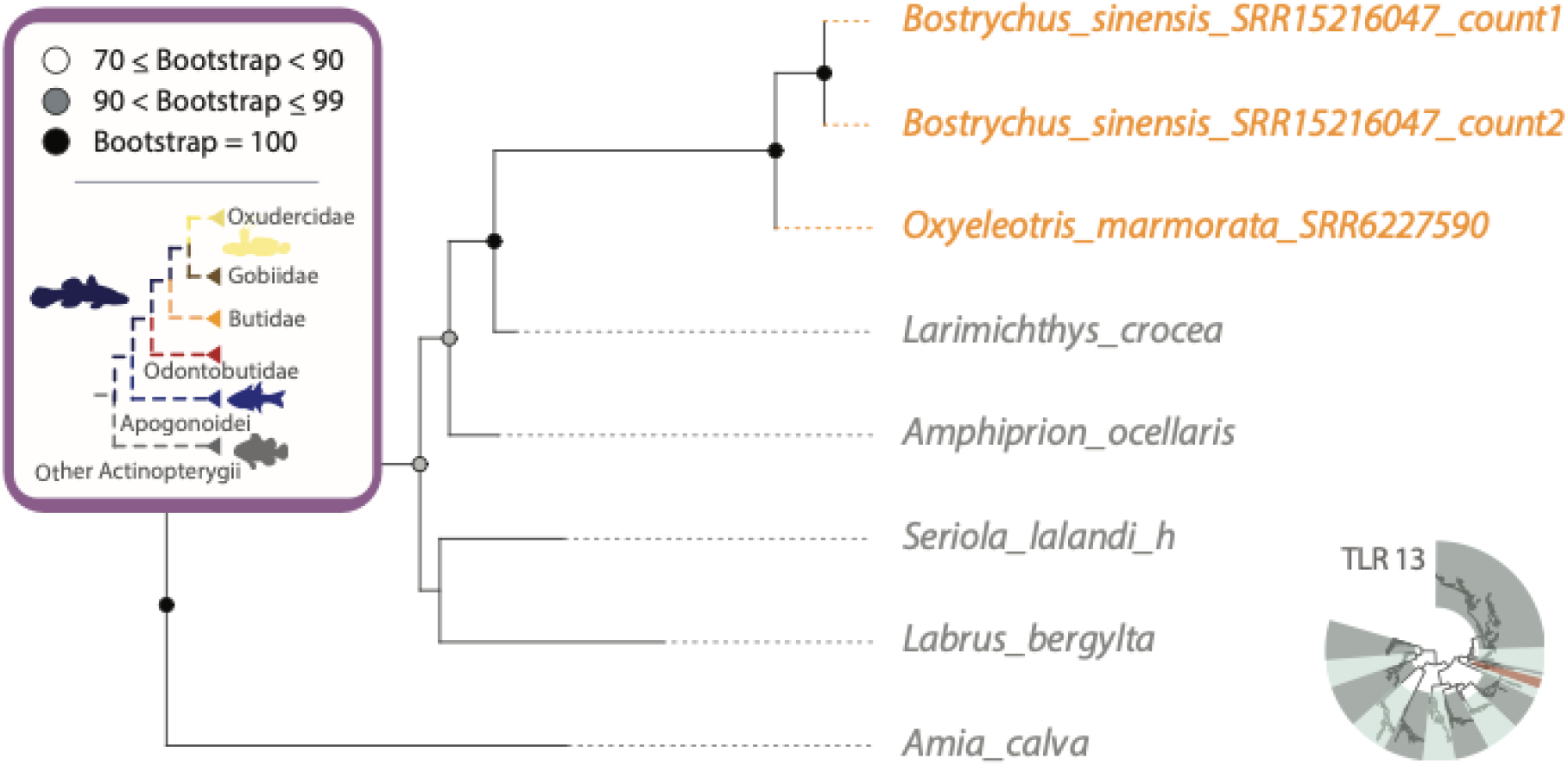
Evolutionary relationships of Gobiiform TLR13 paralogs. Detailed view of the TLR13 phylogeny in **Supplemental Fig. S2**. Oxudercidae (including Mudskippers) are delimited in gold, Gobiidae by brown, Butidae by orange, Odontobutidae by red, and remaining reference actinopterygian taxa by grey. Bootstrap support values 70 or higher are indicated at nodes as indicated in the legend. Sequences without accession numbers were obtained from the TIR dataset (using suggested nomenclature) from Supplemental Table S1 of Carlson et al. (2023). Single letter suffixes in taxon names were used to differentiate taxon names and correspond to the tree file in the data archive.

**Supplemental Fig. S11.**
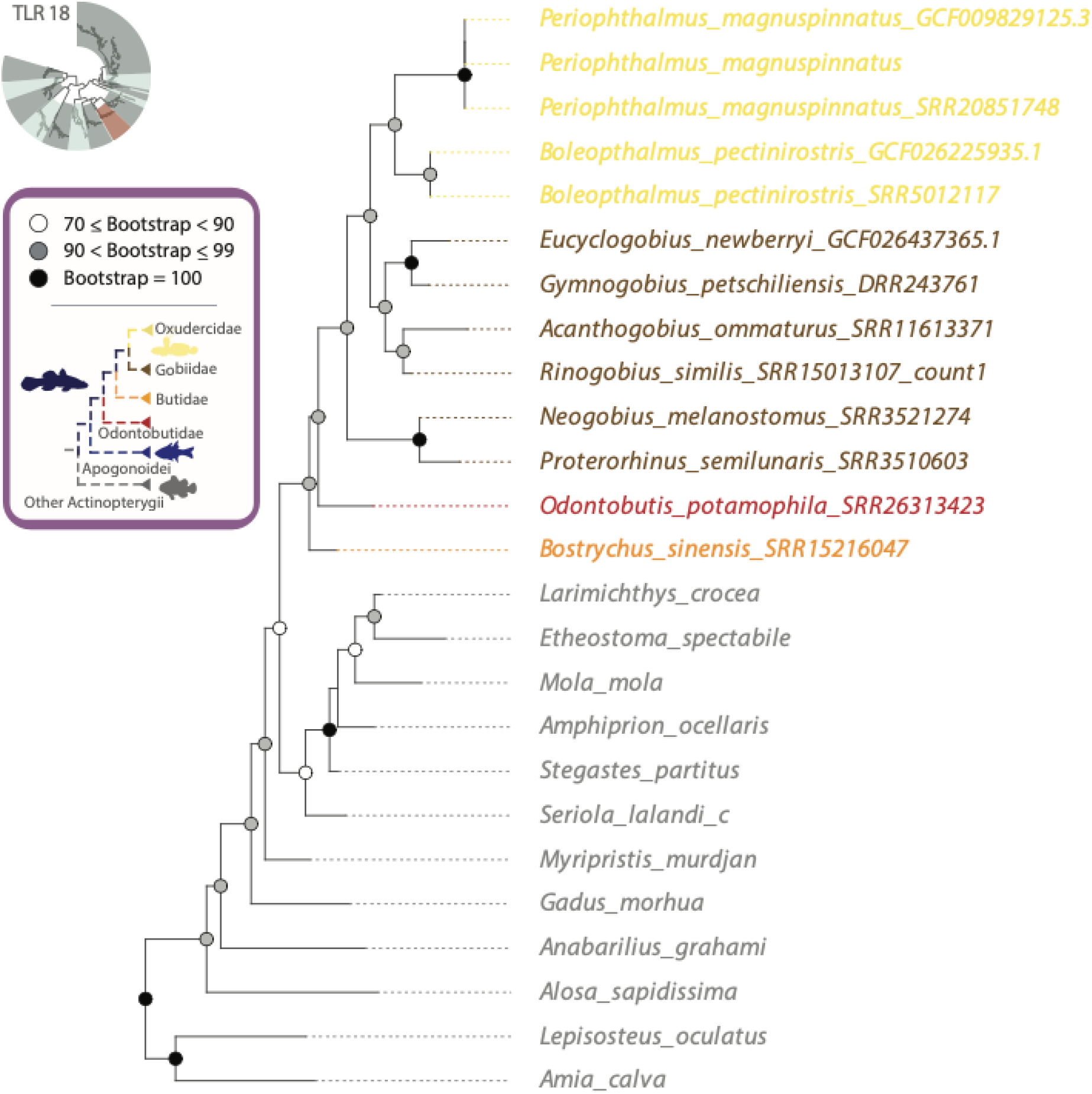
Evolutionary relationships of Gobiiform TLR18 paralogs. Detailed view of the TLR18 phylogeny in **Supplemental Fig. S2**. Oxudercidae (including Mudskippers) are delimited in gold, Gobiidae by brown, Butidae by orange, Odontobutidae by red, and remaining reference actinopterygian taxa by grey. Bootstrap support values 70 or higher are indicated at nodes as indicated in the legend. Sequences without accession numbers were obtained from the TIR dataset (using suggested nomenclature) from Supplemental Table S1 of Carlson et al. (2023). Single letter suffixes in taxon names were used to differentiate taxon names and correspond to the tree file in the data archive.

**Supplemental Fig. S12.**
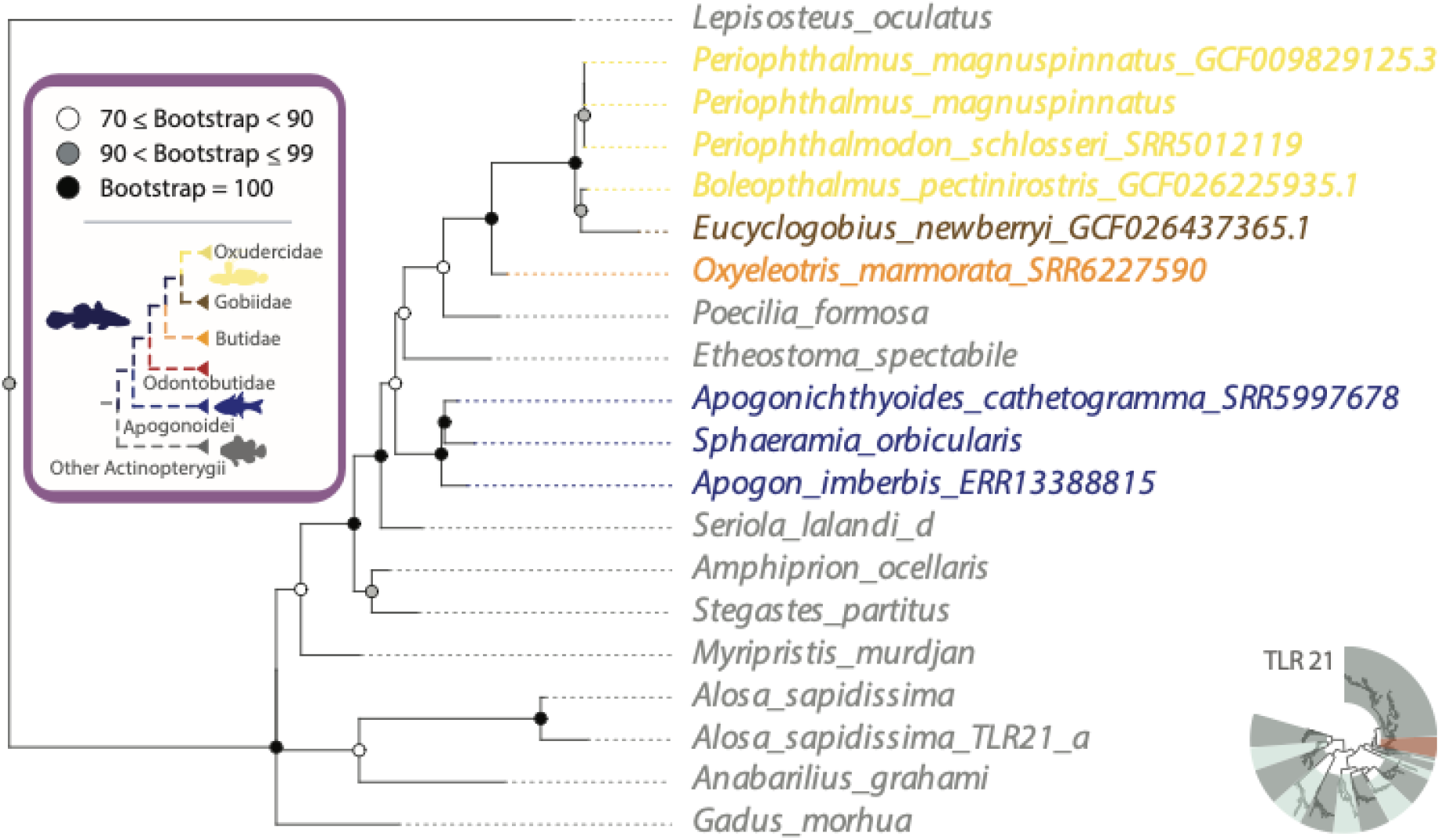
Evolutionary relationships of Gobiiform TLR21 paralogs. Detailed view of the TLR21 phylogeny in **Supplemental Fig. S2**. Oxudercidae (including Mudskippers) are delimited in gold, Gobiidae by brown, Butidae by orange, Odontobutidae by red, and remaining reference actinopterygian taxa by grey. Bootstrap support values 70 or higher are indicated at nodes as indicated in the legend. Sequences without accession numbers were obtained from the TIR dataset (using suggested nomenclature) from Supplemental Table S1 of Carlson et al. (2023). Single letter suffixes in taxon names were used to differentiate taxon names and correspond to the tree file in the data archive.

**Supplemental Fig. S13a.**
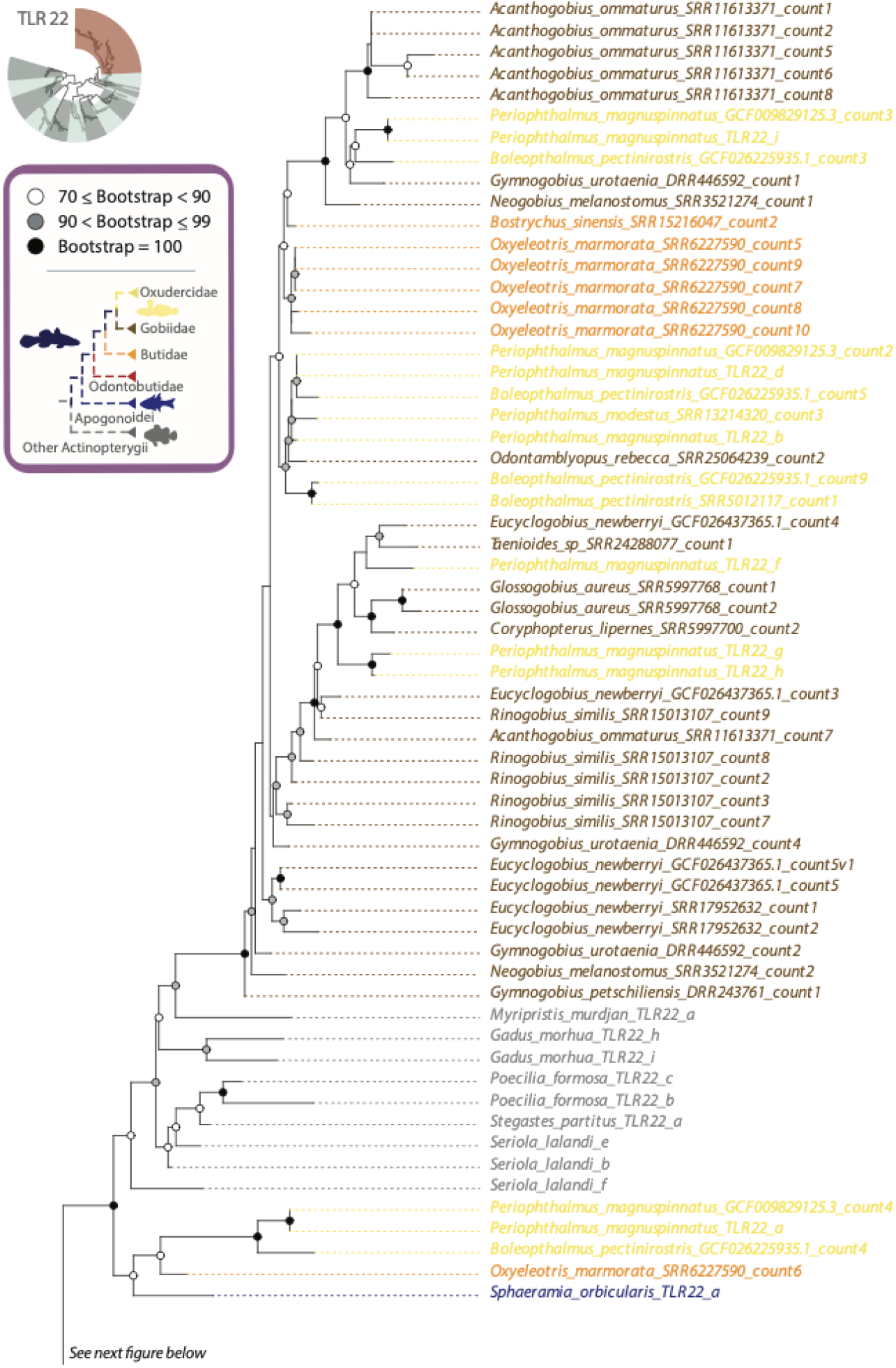
Evolutionary relationships of Gobiiform TLR22 paralogs. Detailed view of the TLR22 phylogeny in **Supplemental Fig. S2**. Oxudercidae (including Mudskippers) are delimited in gold, Gobiidae by brown, Butidae by orange, Odontobutidae by red, and remaining reference actinopterygian taxa by grey. Bootstrap support values 70 or higher are indicated at nodes as indicated in the legend. The sister clade of TLR 22 paralogs is depicted in **Supplemental Fig. S13b**. Sequences without accession numbers were obtained from the TIR dataset (using suggested nomenclature) from Supplemental Table S1 of Carlson et al. (2023). Single letter suffixes in taxon names were used to differentiate taxon names and correspond to the tree file in the data archive.

**Supplemental Fig. S13b.**
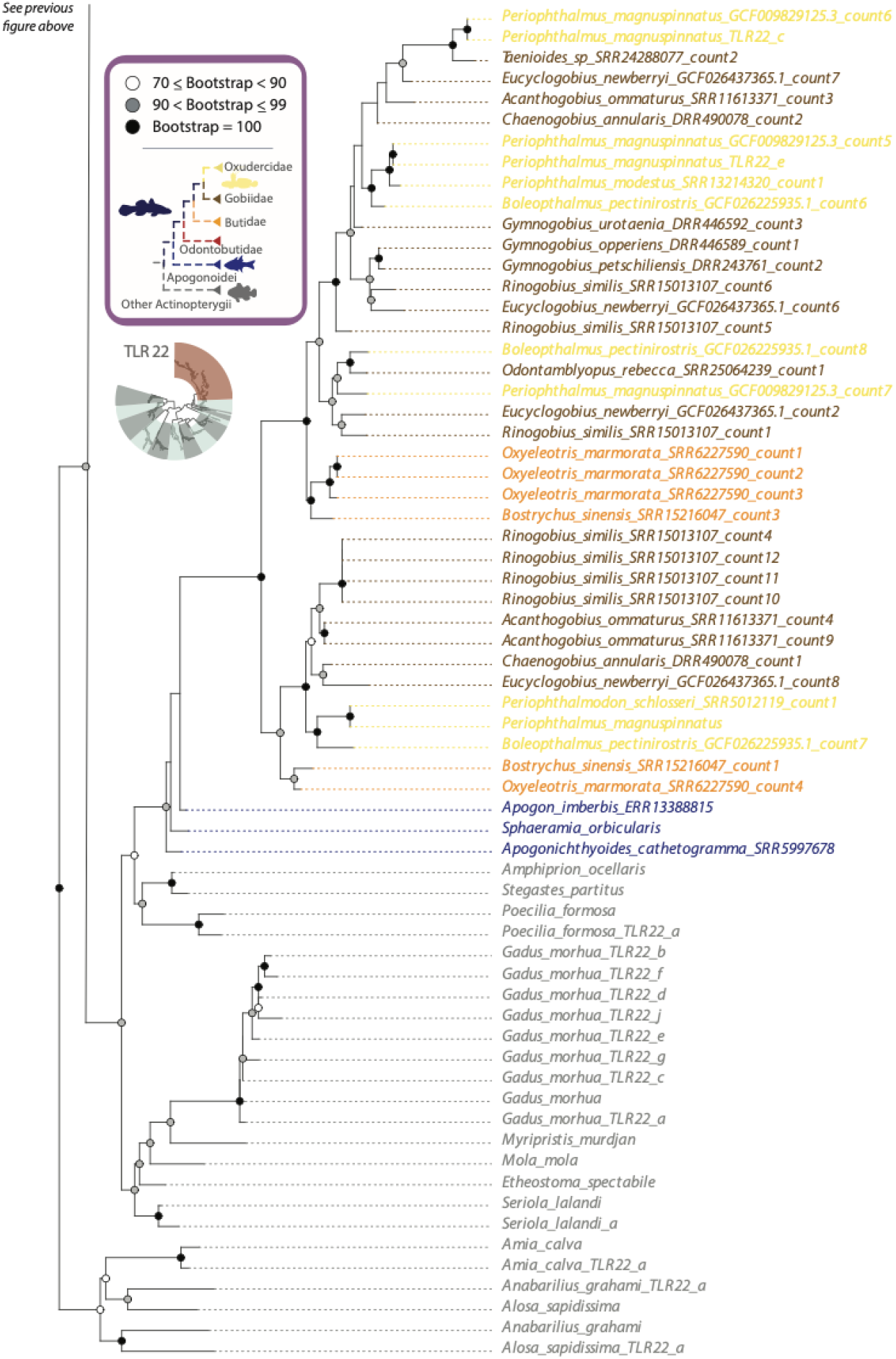
Evolutionary relationships of Gobiiform TLR22 paralogs. Detailed view of the TLR22 phylogeny in **Supplemental Fig. S2**. Oxudercidae (including Mudskippers) are delimited in gold, Gobiidae by brown, Butidae by orange, Odontobutidae by red, and remaining reference actinopterygian taxa by grey. Bootstrap support values 70 or higher are indicated at nodes as indicated in the legend. The sister clade of TLR 22 paralogs is depicted in **Supplemental Fig. S13a**. Sequences without accession numbers were obtained from the TIR dataset (using suggested nomenclature) from Supplemental Table S1 of Carlson et al. (2023). Single letter suffixes in taxon names were used to differentiate taxon names and correspond to the tree file in the data archive.

**Supplemental Fig. S14.**
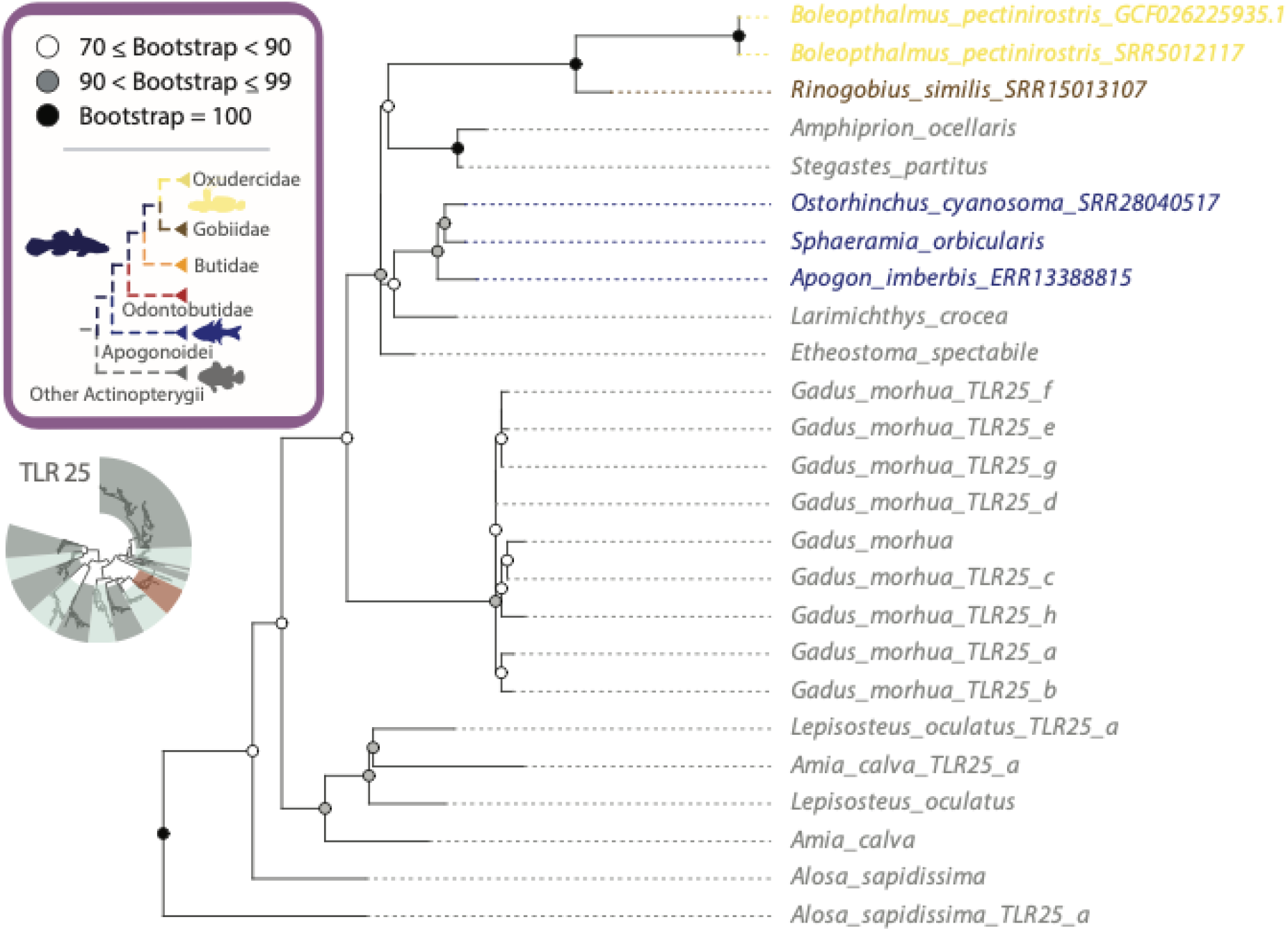
Evolutionary relationships of Gobiiform TLR25 paralogs. Detailed view of the TLR25 phylogeny in **Supplemental Fig. S2**. Oxudercidae (including Mudskippers) are delimited in gold, Gobiidae by brown, Butidae by orange, Odontobutidae by red, and remaining reference actinopterygian taxa by grey. Bootstrap support values 70 or higher are indicated at nodes as indicated in the legend. Sequences without accession numbers were obtained from the TIR dataset (using suggested nomenclature) from Supplemental Table S1 of Carlson et al. (2023). Single letter suffixes in taxon names were used to differentiate taxon names and correspond to the tree file in the data archive.

**Supplemental Fig. S15.**
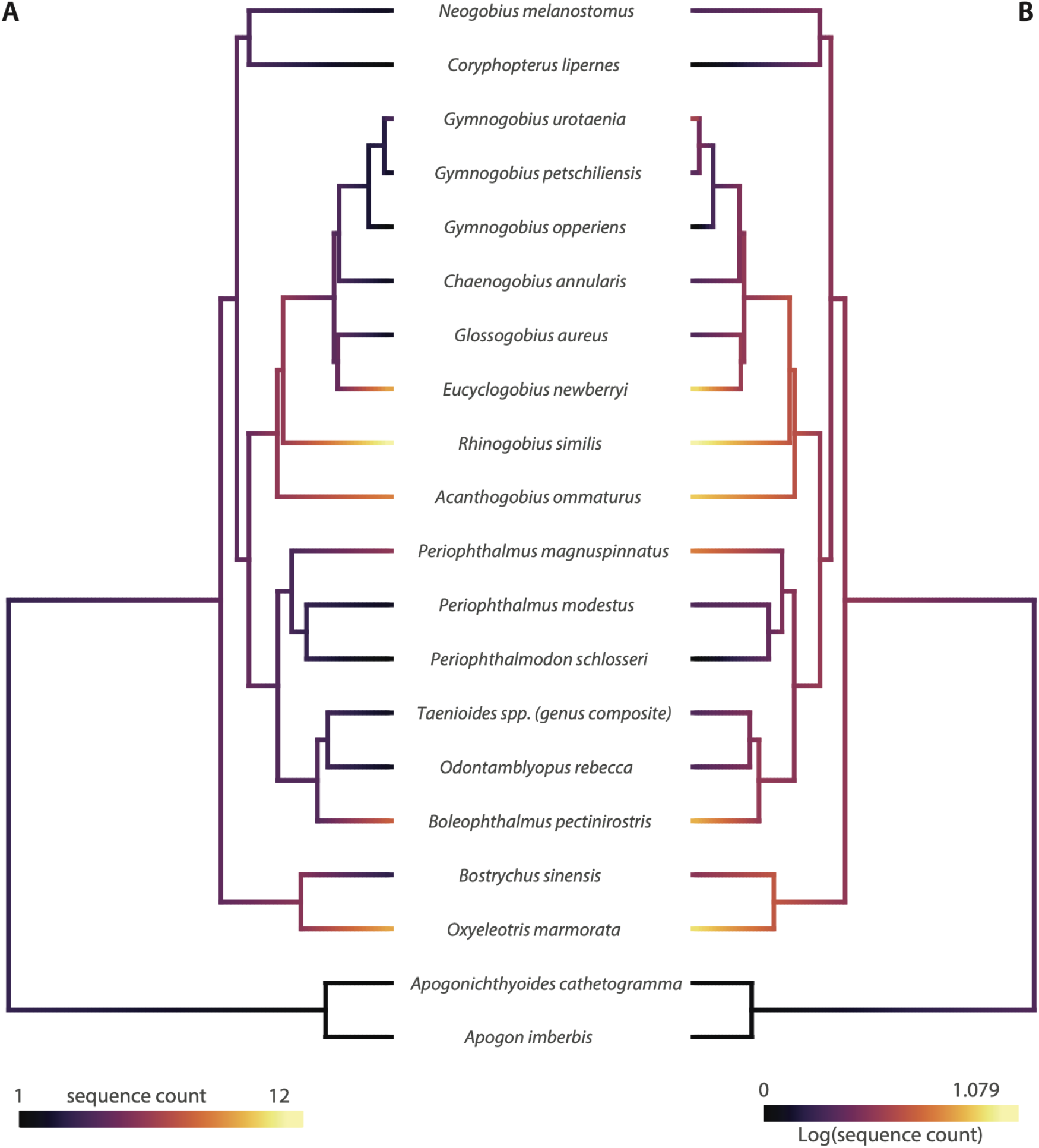
Ancestral State Estimates of TLR22 copy number. Ancestral state estimates of TLR22 copy number variation across the Gobiiform time tree based on raw counts **(A)** or log transformed counts **(B)**. Ancestral state estimates indicated by shadings on phylogenetic branches, with shading intensity corresponding to copy number.

**Supplemental Fig. S16.**
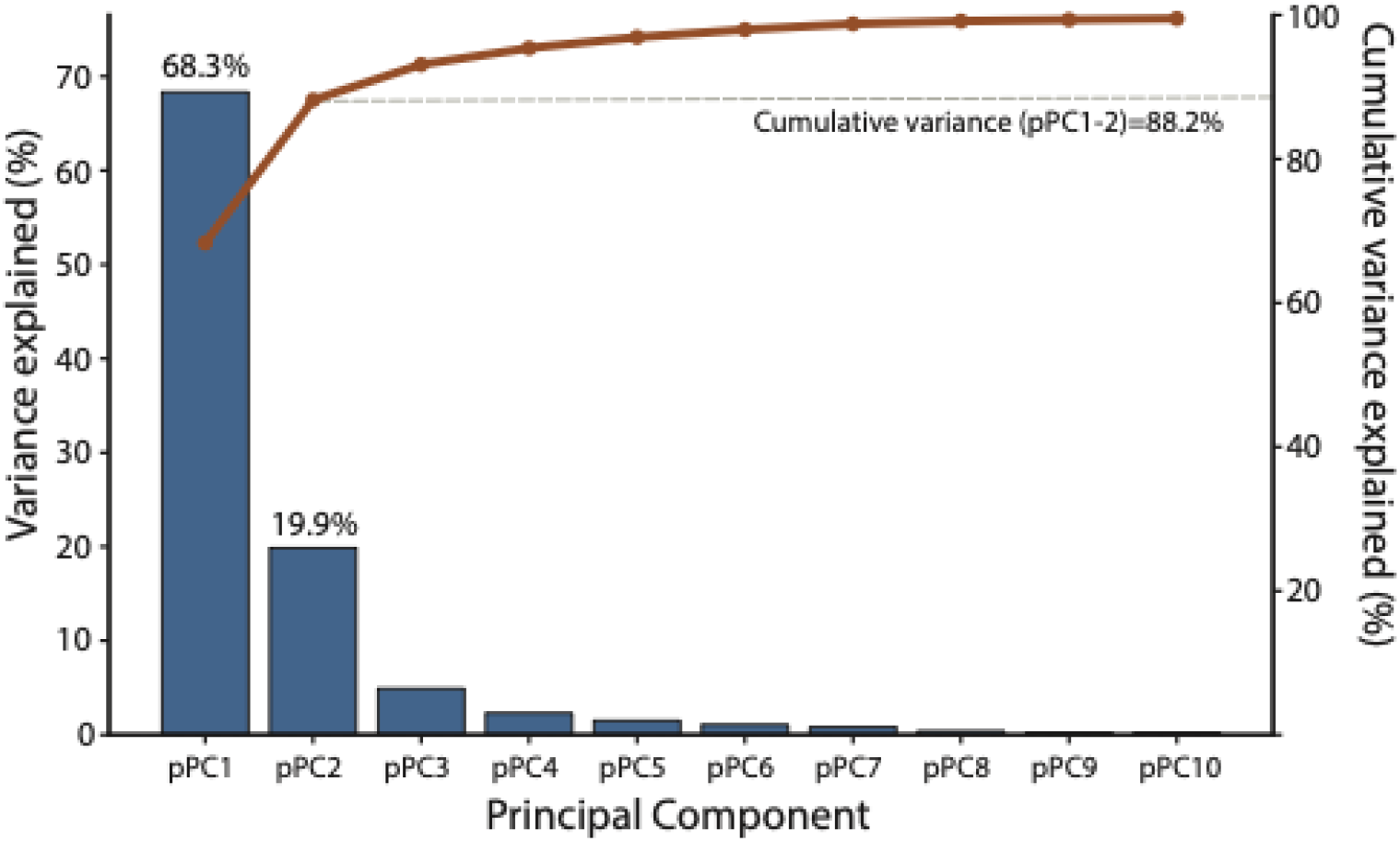
Variance explained by principal components describing TLR protein structural variation. Bars indicate the percentage of total variance explained by each phylogenetic principal component (pPC), and the line indicates cumulative variance explained across successive components. pPC1 and pPC2 explained 68.3% and 19.9% of the total variance, respectively, accounting for 88.2% cumulatively (dashed line). The first ten principal components are shown.

### Supplemental Tables

**Supplemental Table S1:** Habitat data taken from Fishbase (Pauly 2010)

| Scientific name | Marine | Brackish | Freshwater | Aquatic | Amphibious |
| --- | --- | --- | --- | --- | --- |
| *Acanthogobius ommaturus* | --- | TRUE | TRUE | TRUE | --- |
| *Apogon imberbis* | TRUE | --- | --- | TRUE | --- |
| *Apogonichthyoides cathetogramma* | TRUE | --- | --- | TRUE | --- |
| *Boleophthalmus pectinirostris* | TRUE | TRUE | TRUE | --- | TRUE |
| *Bostrychus sinensis* | TRUE | TRUE | TRUE | TRUE | --- |
| *Chaenogobius annularis* | --- | --- | TRUE | TRUE | --- |
| *Cheilodipterus quinquelineatus* | TRUE | --- | --- | TRUE | --- |
| *Coryphopterus lipernes* | TRUE | --- | --- | TRUE | --- |
| *Eucyclogobius newberryi* | TRUE | TRUE | --- | TRUE | --- |
| *Exyrias puntang* | TRUE | TRUE | --- | TRUE | --- |
| *Glossogobius aureus* | --- | TRUE | TRUE | TRUE | --- |
| *Gymnogobius breunigii* | --- | TRUE | TRUE | TRUE | --- |
| *Gymnogobius castaneus* | TRUE | TRUE | TRUE | TRUE | --- |
| *Gymnogobius cylindricus* | TRUE | TRUE | --- | TRUE | --- |
| *Gymnogobius heptacanthus* | TRUE | --- | --- | TRUE | --- |
| *Gymnogobius isaza* | --- | --- | TRUE | TRUE | --- |
| *Gymnogobius macrognathos* | --- | TRUE | --- | TRUE | --- |
| *Gymnogobius mororanus* | TRUE | TRUE | --- | TRUE | --- |
| *Gymnogobius operiens* | TRUE | TRUE | TRUE | TRUE | --- |
| *Gymnogobius petschiliensis* | TRUE | TRUE | TRUE | TRUE | --- |
| *Gymnogobius scrobiculatus* | --- | --- | TRUE | TRUE | --- |
| *Gymnogobius taranetzi* | TRUE | TRUE | TRUE | TRUE | --- |
| *Gymnogobius urotaenia* | TRUE | TRUE | TRUE | TRUE | --- |
| *Neogobius melanostomus* | TRUE | TRUE | TRUE | TRUE | --- |
| *Odontamblyopus rebecca* | TRUE | TRUE | --- | TRUE | --- |
| *Odontobutis potamophila* | --- | --- | TRUE | TRUE | --- |
| *Ostorhinchus cyanosoma* | TRUE | --- | --- | TRUE | --- |
| *Ostorhinchus doederleini* | TRUE | --- | --- | TRUE | --- |
| *Oxyeleotris marmorata* | --- | TRUE | TRUE | TRUE | --- |
| *Paratrypauchen microcephalus* | TRUE | TRUE | --- | TRUE | --- |
| *Periophthalmodon schlosseri* | TRUE | TRUE | TRUE | --- | TRUE |
| *Periophthalmus magnuspinnatus* | --- | --- | TRUE | --- | TRUE |
| *Periophthalmus modestus* | TRUE | TRUE | TRUE | --- | TRUE |
| *Proterorhinus semilunaris* | --- | TRUE | TRUE | TRUE | --- |
| *Rinogobius similis* | --- | --- | TRUE | TRUE | --- |
| *Taenioides sp* | TRUE | TRUE | TRUE | TRUE | --- |

**Supplemental Table S2:** Accessions for individual loci augmenting the ultraconserved element (UCE) dataset.

| Species | COI accession | 12S accession | CYTB accession | RAG1 accession |
| --- | --- | --- | --- | --- |
| *Acanthogobius hasta* | PP933931.1 |  |  |  |
| *Acanthogobius lactipes* |  |  |  | KF415687.1 |
| *Apogon cathetogrammus* | AB890024.1 |  |  | AB893380.1 |
| *Apogon doederleini* | EU381023.1 |  |  | AB893417.1 |
| *Apogon imberbis* | KY176390.1 |  |  | AB893368.1 |
| *Boleophthalmus pectinirostris* | JN033333.1 |  |  | KF415710.1 |
| *Bostrychus sinensis* | OQ553062.1 |  |  |  |
| *Bostrychus zonatus* |  |  |  | KF415713.1 |
| *Chaenogobius annularis* |  |  |  | KF415719.1 |
| *Chaenogobius gulosus* | JX679029.1 |  |  |  |
| *Cheilodipterus quinquelineatus* | MN733703.1 |  |  | JX189812.1 |
| *Coryphopterus lipernes* | GQ367309.1 |  |  | MF049183.1 |
| *Eucyclogobius newberryi* | MF038886.1 |  |  | KF415751.1 |
| *Exyrias puntang* | LC867253.1 |  |  | KF415758.1 |
| *Glossogobius aureus* | PQ637342.1 |  |  | KF415766.1 |
| *Gymnogobius breunigii* | LC867251.1 | KM030451.1 | KF415593.1 | KF415796.1 |
| *Gymnogobius castaneus* | JF952698.1 | LC750854.1 | KF486472.1 | KF486312.1 |
| *Gymnogobius heptacanthus* |  | LC715460.1 | LC577227.1 |  |
| *Gymnogobius isaza* | LC098552.1 | LC717548.1 | AB560891.1 | AB504165.1 |
| *Gymnogobius macrognathos* | LC864460.1 |  |  |  |
| *Gymnogobius mororanus* |  | KM030452.1 | LC577221.1 |  |
| *Gymnogobius operiens* | LC098557.1 | KM030453.1 | LC859615.1 | KF486314.1 |
| *Gymnogobius petschiliensis* | LC098561.1 | LC742205.1 | LC756978.1 | KF486315.1 |
| *Gymnogobius scrobiculatus* | LC864461.1 | LC630803.1 | LC859614.1 |  |
| *Gymnogobius taranetzi* |  | LC385155.1 | KF486477.1 |  |
| *Gymnogobius urotaenia* | OL674313.1 | LC742213.1 | LC730526.1 | KF415797.1 |
| *Neogobius melanostomus* | HQ909495.1 |  |  |  |
| *Neogobius nigricans* |  |  |  | OL412535.1 |
| *Odontamblyopus lacepedii* | MW388860.1 |  |  |  |
| *Odontamblyopus rebecca* |  |  |  | JX003121.1 |
| *Odontamblyopus sp.* |  |  |  | KF415824.1 |
| *Odontobutis obscura* |  |  |  | KF415825.1 |
| *Odontobutis potamophilus* | MK843731.1 |  |  |  |
| *Ostorhinchus cyanosoma* | AB890060.1 |  |  | AB893415.1 |
| *Oxyeleotris marmorata* | KT022088.1 |  |  | KF415829.1 |
| *Paratrypauchen microcephalus* | PP354541.1 |  |  |  |
| *Periophthalmodon schlosseri* | OR053782.1 |  |  |  |
| *Periophthalmus magnuspinnatus* | MW388863.1 |  |  | KP966226.1 |
| *Periophthalmus modestus* | JN033349.1 |  |  | JX003122.1 |
| *Proterorhinus semilunaris* | HQ909494.1 |  |  |  |
| *Rhinogobius flumineus* |  |  |  | KF415857.1 |
| *Rhinogobius similis* | LC867252.1 |  |  |  |
| *Taenioides cirratus* | OM403297.1 |  |  |  |
| *Taenioides sp.* |  |  |  | KF415874.1 |

**Supplemental Table S3:** Calibrations used for RelTime.

| Lower | Upper | Mean | Tip1<sup>1</sup> | Tip2<sup>1</sup> |
| --- | --- | --- | --- | --- |
| 75.4 | 96.5 | 85.9 | *Neogobius sp* | *Kurtus gulliveri* |
| 72.6 | 93.2 | 82.7 | *Kurtus gulliveri* | *Ostorhinchus doederleini* |
| 18.2 | 35 | 26 | *Kurtus gulliveri* | *Kurtus indicus* |
| 55.4 | 80.8 | 68.4 | *Pseudamia gelatinosa* | *Ostorhinchus doederleini* |
| 8.7 | 12 | 10.3 | *Ostorhinchus doederleini* | *Apogon imberbis* |
| 61.3 | 78.7 | 70 | *Rhyacichthys aspro* | *Neogobius sp* |
| 12.5 | 21.7 | 16.9 | *Odontobutis obscura* | *Sineleotris saccharae* |
| 23.6 | 48 | 35 | *Milyeringa veritas* | *Typhleotris madagascariensis* |
| 29.8 | 39.9 | 34.7 | *Ratsirakia lengendrei* | *Erotelis smaragdus* |
| 7.3 | 12.7 | 10 | *Xenisthmus sp* | *Xenisthmus polyzonatus* |
| 16.5 | 23.5 | 19.8 | *Oxyeleotris marmorata* | *Incara multisquamata* |
| 5.4 | 9.5 | 7.5 | *Thalasseleotris iota* | *Grahamichthys radiata* |
| 35 | 45.7 | 40.2 | *Evorthodus minutus* | *Gymnogobius taranetzi* |
| 15.5 | 23.1 | 19.4 | *Gnatholepis scapulostigma* | *Evorthodus minutus* |
| 19.2 | 28.4 | 23.8 | *Periophthalmus magnuspinnatus* | *Boleophthalmus pectinirostris* |
| 8.5 | 14.1 | 11.1 | *Pomatoschistus microps* | *Pomatoschistus minutus* |
| 25 | 33.9 | 29.4 | *Gobiopterus semivestitus* | *Mugilogobius chulae* |
| 36.1 | 46.6 | 41.2 | *Coryphopterus lipernes* | *Neogobius sp* |
| 25.7 | 35.4 | 30.5 | *Glossogobius flavipinnis* | *Bathygobius soporator* |
| 16.7 | 25 | 20.8 | *Kraemeria cunicularia* | *Schismatogobius ninja* |
| 5.8 | 10.1 | 7.8 | *Exyrias bellisimus* | *Istigobius rigillis* |
| 9.2 | 15.1 | 12.1 | *Cryptocentrus cinctus* | *Discordipinna griessingeri* |
| 22.7 | 31.7 | 27.1 | *Bryaninops yongei* | *Eviota sigilata* |
| 10.8 | 19 | 14.8 | *Coryphopterus glaucofraenum* | *Rhinogobiops nicholsii* |
| 13.4 | 22.6 | 17.9 | *Asterropteryx ensifera* | *Vanderhorstia sp* |
| 16.6 | 29.2 | 22.8 | *Callogobius sclateri* | *Callogobius tanegasimae* |
| 21.4 | 30.1 | 25.6 | *Austrolethops wardi* | *Parioglossus dotui* |
| 12.3 | 19.8 | 16 | *Lesueurigobius friesii* | *Lesueurigobius sanzi* |
| 13.4 | 20.7 | 17.2 | *Valenciennea strigata* | *Amblygobius phaelena* |
| 16.7 | 24.8 | 20.6 | *Gobiosoma oceanops* | *Nes longus* |
| 24.9 | 33.5 | 29.1 | *Priolepis cincta* | *Lythrypnus dalli* |
| 15.3 | 23.2 | 19.2 | *Neogobius sp* | *Coryogalops anomolus* |
| 43.2 | 55.6 | 49.2 | *Evorthodus minutus* | *Neogobius sp* |

<sup>1</sup>Tip1 and Tip2 refer to taxa that span the most recent common ancestor of the node calibrated in RelTime (Tamura et al. 2012) conditioned on the ages from McCraney et al. (2025).

